# Cross-cue reconstruction of perceived 3D object structure from human visual cortex

**DOI:** 10.64898/2026.06.08.730830

**Authors:** Shuntaro C. Aoki, Reo Tsukasa, Shiyun Yang, Misato Tanaka, Eizaburo Doi, Tomohiro Nakamura, Jun-Kai Ho, Yukiyasu Kamitani

## Abstract

The human brain assembles three-dimensional (3D) percepts from qualitatively different depth cues, yet the perceived 3D structure that the brain builds—a representation shared across cues—has remained difficult to measure directly. Here, we show that this cue-invariant 3D structure can be externalized as explicit 3D objects from human brain activity: fMRI responses are decoded into the latent features of a pretrained 3D point-cloud autoencoder, and a generator then maps these features back to a point cloud. A decoder trained exclusively on responses to 2D rendered objects passed three increasingly stringent tests: (i) it generalized to novel object categories; (ii) it generalized across depth cues to random dot stereograms (RDSs), which evoke 3D percepts through binocular disparity but share no pictorial shape information with the training images; and (iii) it tracked the 3D slant of contour-matched RDSs whose 2D outlines were held identical but whose disparity-defined slants varied, indicating that the reconstruction reflected depth-defined geometry rather than object category or image outline. Cross-cue generalization was strongest in higher visual areas, particularly along the dorsal stream. These results indicate that cross-cue generalization can serve as a criterion for externalizing perceived 3D structure and open a route toward reading out internal 3D representations that go beyond the momentary retinal input and could support predictions of how the world would appear under different viewpoints—a step toward externalizing the brain’s internal world model.

## Introduction

The human brain infers the three-dimensional (3D) structure of the external world from visual input arriving as two-dimensional (2D) retinal images. This 2D-to-3D inference depends on depth cues embedded in retinal images^1^. Depending on viewing context, the same perceived 3D structure can arise from one or more qualitatively distinct cues, including monocular pictorial cues (shading^2–4^, textures^5,6^, edge/contour, perspective) and binocular disparity^7^. Each cue, alone or in combination, can evoke a coherent 3D percept. Neurophysiological and neuroimaging studies have also identified depth representations that respond to 3D structure regardless of which depth cue (e.g., shading or binocular disparity) carries that structure on the retina^8–10^.

Together, these observations suggest that the brain constructs a cue-invariant internal representation of 3D structure—a model of the external world that is distinct from retinal inputs and can be assembled from diverse visual inputs sharing the same underlying 3D structure (Figure 1A). Such an internal 3D representation is richer than a copy of the momentary retinal input: it can support predictions about how the same object would appear from another viewpoint and how it would behave under other actions, and in this sense it functions as part of the brain’s internal “world model” of the external environment^11,12^. Yet existing approaches have mainly characterized these representations through similarity analyses of cortical response patterns. Such analyses can detect statistical similarities and differences between responses elicited by different cues, but statistical detectability is not access to the perceived 3D structure itself: whether cortical signals can be translated into the 3D structure that the brain constructs from those cues, rather than only into a statistical claim about it, remains open.

**Figure 1.**
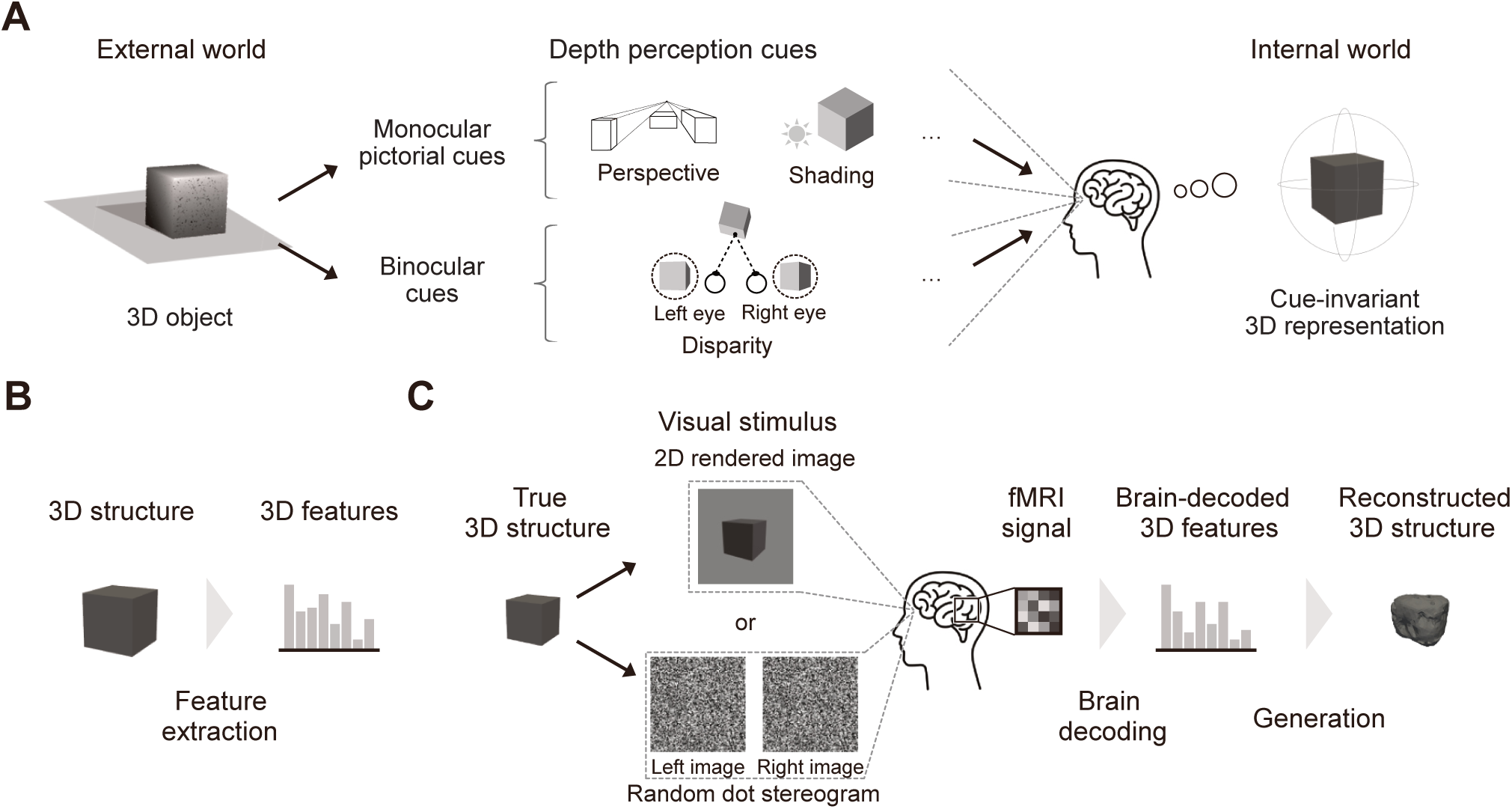
3D reconstruction framework. (A) Brain construction of cue-invariant 3D structure from diverse depth cues. (B) Extraction of 3D latent features from explicit 3D objects with a pretrained point-cloud autoencoder. (C) Cross-cue reconstruction pipeline. Subjects view each object as a 2D rendered image (monocular pictorial cues) or a random dot stereogram (RDS; binocular disparity) during fMRI. A trained brain decoder predicts 3D latent features from fMRI signals, and the autoencoder’s generator converts them into an explicit 3D structure.

Externalizing this representation requires translating brain activity into an explicit 3D object, such as a mesh or point cloud, that can be inspected directly. If the brain constructs a 3D percept of the external world from retinal cues, the externalized content should not depend on which cue evoked the underlying brain activity. Cross-cue reconstruction—reconstructing matching 3D content from brain activity elicited by qualitatively different depth cues—therefore tests whether externalization captures perceived 3D structure rather than cue-specific correlations. Reconstruction of perceptual content from brain activity has advanced substantially in the 2D domain: techniques that leverage deep neural network (DNN) features have enabled the reconstruction of subjective visual experiences as images^13–15^. These studies externalize perceptual content, but 2D image reconstruction remains in the format of retinal input and cannot reveal 3D representations that the brain infers beyond that input.

Prior studies have begun to read out 3D information from brain activity, but they have not established access to a cue-invariant 3D representation. Zheng et al.^16,17^ decoded depth-defined contrast images from functional MRI (fMRI), capturing binocular disparity information without testing generalization across alternative depth cues. Lescroart and Gallant^18^ showed that scene-selective cortical regions represent the 3D configuration of large surfaces and produced coarse reconstructions of scene structure, but their stimuli were limited to 2D rendered images, and the reconstructions were not evaluated across depth cues. Gao et al.^19^ attempted explicit 3D shape reconstruction from brain activity elicited by 360^◦^ object videos; however, the reconstruction relies heavily on 2D visual features and semantic information, and cue-invariant generalizability was not examined. Consequently, it remains unknown whether the reconstructed 3D information in these studies reflects a common, cue-invariant 3D representation or cue-specific correlations.

Here, we ask whether a cue-invariant internal representation of 3D structure can be externalized from human brain activity as an explicit 3D object—a mesh or point cloud. Specifically, we test whether brain activity evoked by visual stimuli that convey depth through different cues (2D rendered images and random dot stereograms, RDSs) can be used to reconstruct the viewer-centered 3D structure that the subject perceives. We refer to this view-dependent 3D structure hereafter as a perceived 3D structure.

We adapt the translator–generator framework^13,14,20^, originally established for 2D image reconstruction from brain activity, to explicit 3D point clouds (Figure 1B, C). As a first step, we restrict the framework to single isolated objects against a blank surround. A DNN trained on 3D instances first encodes each ground-truth perceived 3D structure into latent 3D features, which serve as an intermediate representation. The pipeline then reconstructs a 3D point cloud in two stages (Figure 1C): (i) a translator—the brain decoder—maps fMRI responses to these latent 3D features, and (ii) a generator converts the decoded features back into a 3D point cloud. We hypothesize that latent features learned by such DNNs encode 3D information aligned with the brain’s own 3D representations, allowing neural 3D information to be accessed through the latent space.

We evaluate the pipeline with three independent tests of increasing stringency: (i) reconstruction of perceived 3D structures from novel object categories, (ii) generalization across qualitatively different depth cues (monocular pictorial cues vs. binocular disparity), and (iii) reconstruction from stimuli whose 2D contour is held constant while only the 3D slant varies. Reconstruction across these conditions—especially under the contour-matched test—would support cross-cue generalization as a criterion for externalizing perceived 3D structure and indicate that the pipeline taps a cue-invariant 3D representation in the human brain.

## Results

To probe whether the pipeline exploits the brain’s intrinsic 3D information rather than semantic content or low-level retinal features, we collected fMRI responses from five subjects (S1–S5) while they viewed 2D rendered images of objects (2,000 training stimuli, each presented three times; Figure 2). We then evaluated the trained pipeline across a graded series of test sets (Figure 2): (i) new natural objects from trained categories (baseline), (ii) new natural objects from novel categories plus 30 abstract artificial objects (to rule out semantic shortcuts), (iii) the same artificial objects presented as random dot stereograms (RDSs), which share no pictorial shape information with the training images (to test cross-cue generalization), and (iv) contour-matched RDSs in which the projected 2D contour was identical across stimuli but the underlying 3D slant varied (to rule out reliance on disparity-defined outline). Each test stimulus was presented eight times, and the fMRI responses were averaged across repetitions. Neural activity was extracted from five regions of interest in the visual cortex (VC), defined on the HCP MMP1.0 cortical parcellation^21^: early VC, MT & neighbors, dorsal VC, ventral VC, and the entire VC (whole VC; Table 1). In the following, we first ask whether visual-cortical activity predicts the 3D latent features, then reconstruct explicit 3D structures from these predictions, isolate depth-cue-invariant slant using contour-matched RDSs, and finally compare cortical regions.

**Figure 2.**
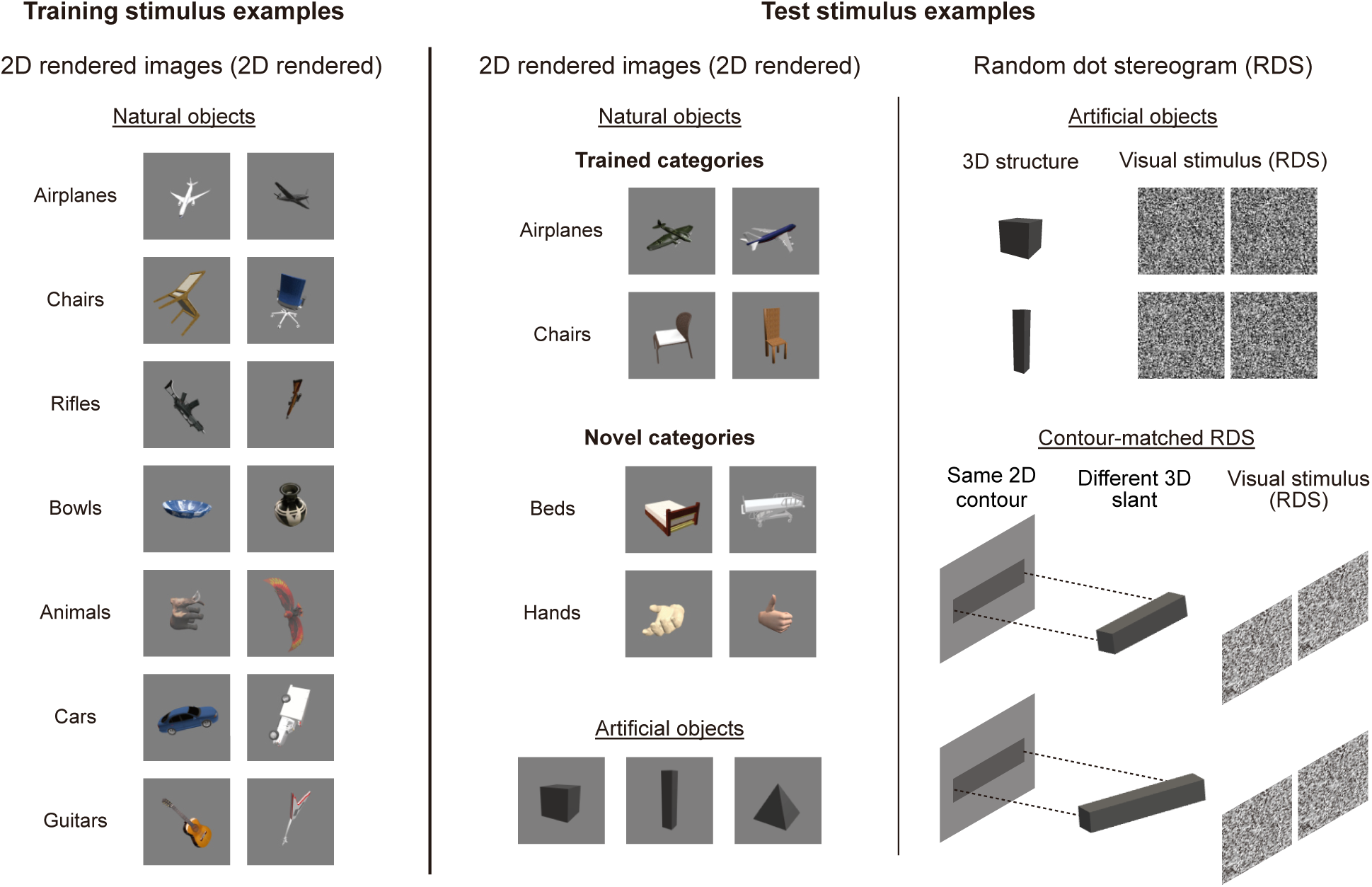
Visual stimuli. Left: 2D rendered training images of natural object categories (e.g., airplanes, chairs, rifles), each from a single viewpoint. Middle: 2D rendered test images: (i) natural objects from trained categories, (ii) natural objects from novel categories, and (iii) artificial objects. Right: Random dot stereogram (RDS) test stimuli: (i) artificial-object RDSs and (ii) contour-matched RDSs in which the projected 2D contour was held constant while the underlying 3D slant varied. All test objects were new exemplars not shown during training; the natural objects are grouped only by whether their object category appeared in the training set (trained vs novel categories).

**Table 1.** Visual cortical ROI definitions. Each region of interest (ROI) in the visual cortex (VC) is the union of the listed HCP-MMP1.0 parcels. The whole VC additionally includes DVT, ProS, POS1, and POS2.

| ROI | HCP-MMP1.0 parcels |
| --- | --- |
| Early VC | V1, V2, V3 |
| MT & neighbors | V3CD, LO1, LO2, LO3, MT, MST, V4t, FST, PH, TPOJ2, TPOJ3 |
| Dorsal VC | V3A, V3B, V6, V6A, V7, IPS1, MIP, LIPv, LIPd, VIP, AIP, 7AL, 7Am, 7PC, 7PL, 7Pm, PGp, IP0, IP1, IP2 |
| Ventral VC | V4, V8, VVC, VMV1, VMV2, VMV3, PIT, FFC, PHA1, PHA2, PHA3 |
| Whole VC | Union of the four ROIs above, plus DVT, ProS, POS1, POS2 |

### Decoding 3D latent features from cortical activity

We first establish that the autoencoder latent features form an intrinsic 3D target, and then test whether they can be decoded from cortical activity. To externalize 3D structures from brain activity, we used the latent features of pretrained 3D point-cloud autoencoders as decoding targets (Figure 1C). We trained two complementary autoencoders on 3D point clouds—without 2D images or semantic labels—and used them as parallel backbones: AtlasNet^22^ (Figure S1A) as the primary architecture for the main results, and a diffusion-based point-cloud autoencoder^23^ as a robustness check. All main figures and tables report results from the AtlasNet autoencoder; corresponding results from the diffusion-based autoencoder are provided in the supplementary figures. We then verified that the resulting latent features possess three key properties expected of an intrinsic 3D representation.

First, latent features yielded near-perfect recovery of the input point clouds (Figure S1B). Second, linear interpolation between two latent features yielded recovered point clouds that morphed smoothly between the endpoints (Figure S1C). We quantified this continuity using the Chamfer distance (see Methods, Chamfer distance; Figure S1D), a symmetric average of nearest-neighbor distances between two point clouds (intuitively, a measure of how completely each cloud is covered by the other). The Chamfer distance between the recovered point clouds increased monotonically along the interpolation trajectory (Figure S1E).

Third, representational similarity analysis (RSA; Figure S2) compared the distance structure of the AtlasNet autoencoder’s latent features with those based on rendered-image pixels, semantic categories, and 3D point clouds. Latent-space distances correlated strongly with the Chamfer distance between 3D point clouds but not with pixel- or semantic-space distances, indicating that the AtlasNet autoencoder relies on 3D information rather than pixel-level image content or semantic category. Together, these results suggest that the AtlasNet autoencoder’s latent features encode intrinsic 3D representations suitable for our pipeline. The same three properties were also observed with the diffusion-based autoencoder (Figure S3), confirming that they generalize beyond AtlasNet.

Based on these results, we asked whether the decoders trained exclusively on fMRI responses to 2D rendered images could predict the DNN latent features of novel objects. For each training image, we extracted the unit activations from the encoder of the 3D point-cloud autoencoder (Figure 3A) and used them as decoding targets. We trained an L2-regularised linear decoder for every latent unit, using the 5,000 voxels within each ROI whose responses were most strongly correlated with that unit across the training set. During testing (Figure 3B), fMRI responses to previously unseen stimuli were fed into the trained decoders to generate decoded latent features.

**Figure 3.**
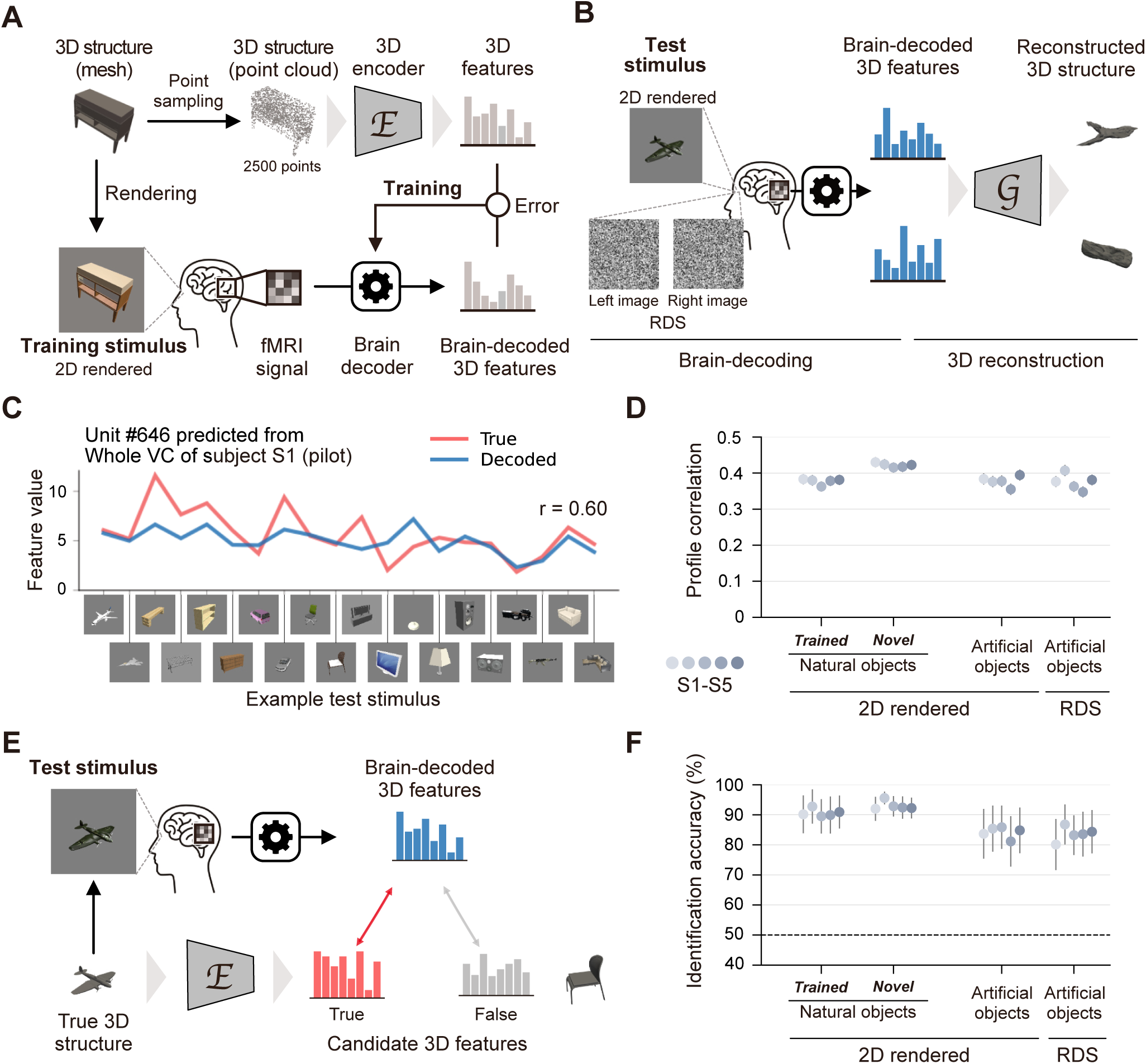
Decoding 3D latent features. (A) Training: subjects viewed 2D rendered images paired with ground-truth latent features from a pretrained point-cloud autoencoder; ROI-wise multivoxel patterns were mapped to the features by L2-regularised linear regression. (B) Testing and reconstruction: trained decoders were applied to novel 2D rendered images and random dot stereograms (RDSs) to predict 3D latent features (brain-decoded features in blue), which the autoencoder’s generator *G* then converted into a reconstructed 3D structure. (C) Example unit (#646, subject S1) across test stimuli; ground-truth feature in red, whole-VC decoded feature in blue. (D) Whole-VC profile correlation (per-unit Pearson’s correlation across stimuli) between decoded and true features, by stimulus set. Each dot is one subject (S1–S5); error bars are 95% CIs. (E) Pairwise identification: for each test stimulus, the decoded feature is compared by Pearson’s correlation with the true and a false candidate; a trial is correct when correlation is larger for the true candidate. (F) Whole-VC identification accuracy by stimulus set. Each dot is one subject (averaged across test stimuli); error bars are 95% CIs; dashed line indicates chance (50%).

Decoding performance was quantified by two complementary measures (see Methods, Decoding evaluation). The first, the *profile correlation*, is the Pearson’s correlation across test 3D structures between a unit’s decoded and ground-truth feature values, capturing how well its response profile over stimuli is recovered (Figure 3C). For the whole VC, the mean profile correlation across units was about 0.40 (Figure 3D), comparable to values reported in a previous 2D feature decoding study^24^. The second is pairwise identification accuracy at the feature level (Figure 3E), which quantifies how accurately the true 3D structure can be identified from a false alternative using the similarity between decoded and candidate latent features. Mean identification accuracy across decoded samples for the whole VC exceeded 90% for natural objects from both trained and novel categories (Figure 3F), and was significantly above chance (50%) in every subject (one-sample *t* test across stimuli).

Identification accuracy for artificial objects under RDSs also exceeded 80%, comparable to that under 2D rendered images (Figure 3F). Because the decoders were trained exclusively on 2D rendered images containing pictorial depth cues, this comparable performance under RDSs—which convey 3D through binocular disparity and share no pictorial shape information with the training set—is a first indication that the decoders generalize across qualitatively different depth cues at the feature level.

Comparable identification performance was obtained with the latent features from the diffusion-based autoencoder, although the profile correlations were slightly lower (Figure S3E), suggesting that these results are not specific to a particular network architecture. Together, these results show that 3D latent features can be predicted reliably from human visual-cortical responses by decoders trained solely on fMRI responses to 2D rendered images, and that the decoders generalize to fMRI responses elicited by different depth cues. The reliable prediction suggests that the autoencoder latent features are aligned with visual-cortical representations of 3D structure, whereas the generalization from 2D rendered images to RDSs further indicates that this aligned representation is not tied to the pictorial cues present in the training images but includes a cue-invariant component.

### Reconstructing 3D structures from brain activity

Having established that 3D latent features can be decoded from fMRI activity, we next reconstructed explicit 3D structures with the two-stage translator-generator framework (Figure 3B). The decoder first translates multivoxel brain activity into 3D latent features; the autoencoder’s generator then converts them into a 3D point cloud.

Whole-VC responses to 2D rendered images yielded recognisable 3D reconstructions for natural objects in Figure 4A and artificial objects in Figure 4B; comparable reconstructions were obtained with the diffusion-based autoencoder (Figure S4A), confirming that the results are not specific to a particular network architecture. Additional examples are provided in Figures 4C, 4D, S5A, and S5B. The reconstructions reproduced the coarse structure of the true 3D structures, including overall shape and orientation in 3D space. In some cases, they captured diagnostic parts such as the back and legs of a chair, although fine structural details, such as the tail of an airplane, tended to be lost. The rotating view confirmed that the pipeline reconstructed full 3D structure rather than a single 2D projection. Reconstructions varied considerably within each category (Figures 4C and S5A); the quantitative test against a category-prototype account is reported below. Reconstruction quality was qualitatively comparable for objects from training and novel categories, underscoring the generalizability of both the feature decoders and the reconstruction pipeline.

**Figure 4.**
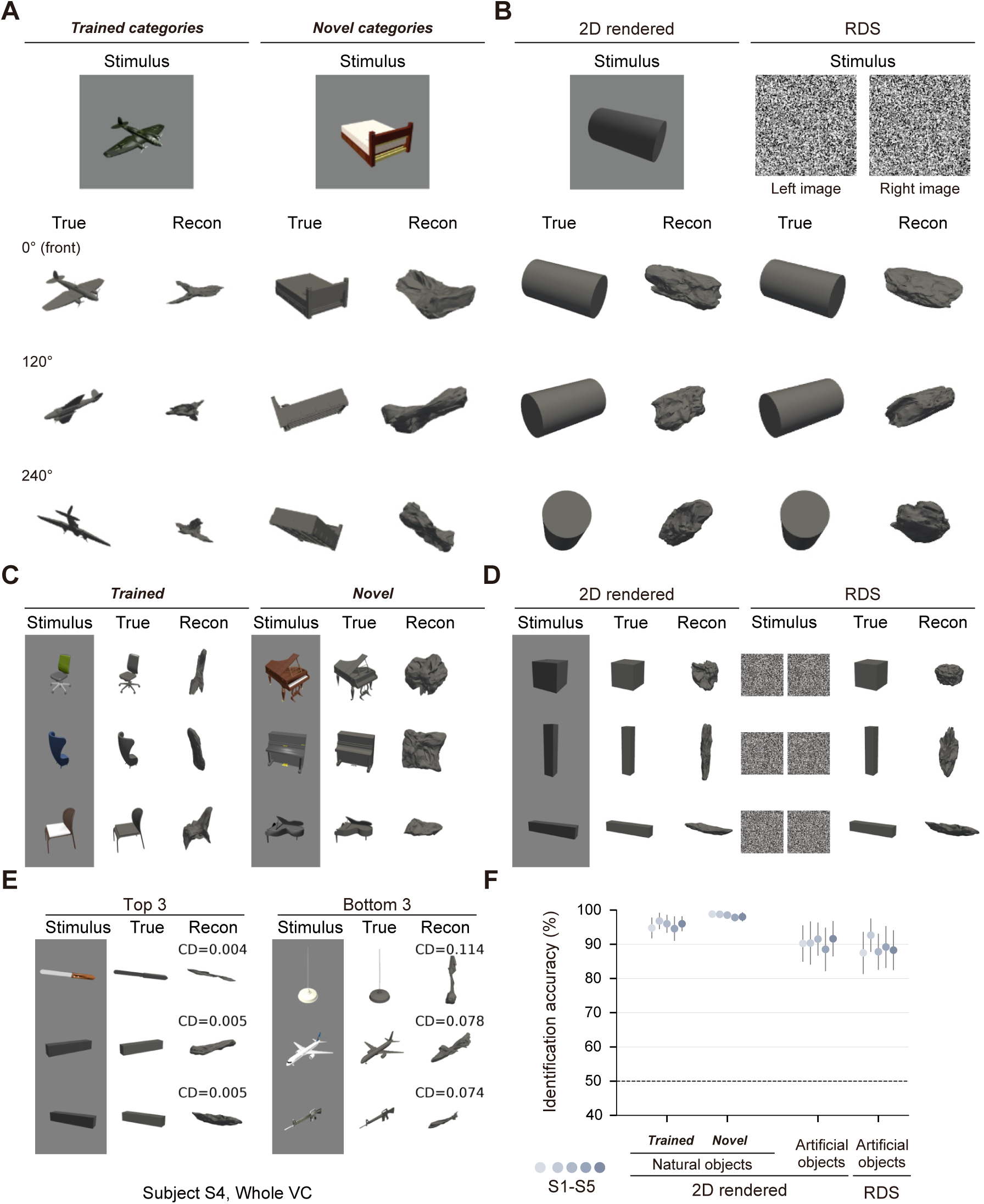
3D reconstruction. (A) Example whole-VC reconstructions of natural objects (2D rendered images) for subject S4—one trained-category and one novel-category object—each shown as the stimulus together with the ground-truth (True) and reconstructed (Recon) 3D structures from three viewpoints (0^◦^, 120^◦^, 240^◦^). (B) Example whole-VC reconstructions of an artificial object under 2D rendered images and RDSs, in the same format. (C) Additional natural-object reconstructions grouped by trained and novel categories, illustrating within-category variability of the reconstructions. (D) Additional artificial-object reconstructions under 2D rendered images and RDSs. (E) Best (left, Top 3) and worst (right, Bottom 3) reconstructions of trained-category natural objects, ranked by Chamfer distance (CD). (F) Identification accuracy by stimulus set. Each dot is one subject (S1–S5); error bars are 95% CIs; dashed line indicates chance (50%). Rotating-video reconstructions are available online (natural objects: https://www.youtube.com/watch?v=RGrhT6gomMo; artificial objects, 2D rendered and RDSs: https://www.youtube.com/watch?v=YzQQ4tl3_oM).

Reconstructions of comparable quality were also obtained when artificial objects were presented as random dot stereograms (RDSs), although the decoders were trained exclusively on fMRI responses to 2D rendered images containing pictorial depth cues (Figure 4B, RDS columns; Figures 4D and S5B). This indicates that the feature decoders and the reconstruction pipeline capture aspects of 3D structure that generalize across qualitatively different depth cues.

Reconstructions were also highly consistent across subjects, sharing the coarse shape and orientation of the true 3D structures (Figure S6A).

Chamfer distance quantified how closely each reconstruction matched its ground-truth 3D structure in global point-to-point coverage (Figure S1D). Among natural objects from trained categories, the three best and three worst reconstructions ranked by Chamfer distance are shown in Figure 4E. Coarse object shape and overall orientation were captured well in the best examples, especially for elongated objects, whereas the worst reconstructions lost key structural elements and fine-scale detail. Nevertheless, even in these poorer cases, global orientation was often preserved, indicating that larger Chamfer distances do not necessarily imply complete re-construction failure. The Chamfer distances for each reconstructed object were highly correlated between S1 and each of the other subjects, aligning with the cross-subject consistency observed by visual inspection (Figure S6B).

We complemented the quantitative evaluation by an identification analysis based on Chamfer distance. Each reconstructed structure served as a probe and was compared with its true counterpart and false alternatives drawn from the corresponding candidate set of 3D structures (natural or artificial objects); a trial was counted as correct when the Chamfer distance to the true 3D structure was smaller than to the false one. For 2D rendered images, identification accuracy exceeded 90% for natural objects from both trained and novel categories and remained high (*>* 80%) for artificial objects (Figure 4F; above chance in every subject). Moreover, in a more stringent same-category pairwise identification analysis in which false candidates were restricted to 3D structures from the same category as the true object (chance = 50%), accuracy still exceeded 90% on average for both trained and novel categories (Figure S7A), ruling out the alternative explanation that the pipeline merely decodes category-level information rather than object-specific 3D structure.

For artificial objects presented as RDSs, identification accuracy also exceeded 80% (Figure 4F). Because the decoders were trained exclusively on 2D rendered images containing pictorial depth cues, this comparable RDS performance supports generalization across qualitatively different depth cues at the reconstruction level. A group-level linear mixed-effects model (LMM) analysis revealed no significant difference in identification accuracy between the two stimulus formats (*β*_RDS−rendered_ = −1.4%, 95% CI [−6.2, +3.5], two-sided *p* = 0.477; the Bayesian posterior was consistent: Pr(dir.) = 0.744; Tables S1–S2; see Methods, Statistical analyses, for the group-level inference framework).

Comparable patterns were observed with the diffusion-based autoencoder (Figure S4B), indicating that reconstruction success does not depend on the particular DNN. Two additional checks reinforced these findings: identification accuracy based on human pairwise judgments closely tracked the Chamfer-distance-based accuracy (Figure S7C), and reconstruction quality remained above 80% even when decoding from noisier single-trial responses (Figure S7B).

Together, these results show that visual cortical activity supports reconstruction of explicit 3D structures that preserve the coarse shape and 3D orientation of the underlying objects. The same reconstruction also generalized from 2D rendered images to RDSs, indicating that the decoder accesses information consistent with cue-invariant 3D representations.

### Predicting slant from contour-matched random dot stereograms

The RDS reconstructions still left a possible non-3D confound: each artificial-object RDS had a unique disparity-defined contour that could identify the object. The pipeline might therefore exploit 2D contour identity rather than 3D information. To rule out this confound, we designed a second RDS stimulus set in which the projected 2D contour was held constant while the underlying 3D structures varied (Figure 5A). Specifically, bar-like objects were slanted to multiple orientations in depth, and each bar was deformed so that its projection onto the fronto-parallel plane—the cyclopean silhouette, not the individual retinal images—produced an identical outline. Consequently, the stimuli shared the same cyclopean 2D outline while their binocular disparity, and hence 3D slant, varied, providing a stricter test of cross-cue reconstruction. We then asked whether the pipeline could still reconstruct slant in 3D space when observers viewed these contour-matched RDSs.

**Figure 5.**
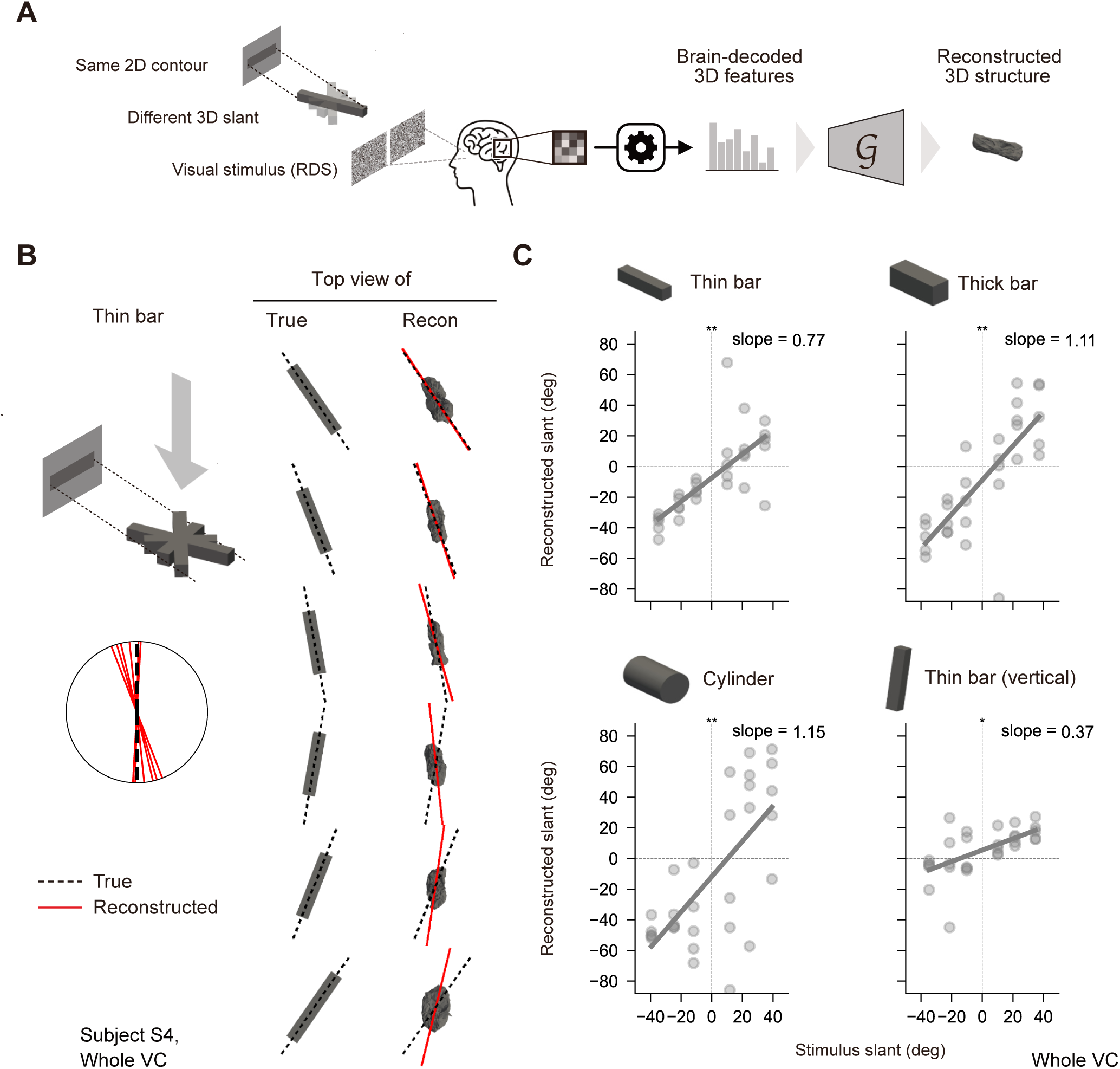
Slant prediction from contour-matched RDSs. (A) Contour-matched RDSs share an identical cyclopean 2D contour (fronto-parallel projection), not matched retinal images, but differ in disparity-defined 3D slant; whole-VC fMRI patterns are decoded and reconstructed. (B) Example whole-VC reconstructions of a horizontal bar across stimulus slants (top views), shown enlarged for clarity. Red solid / black dashed lines: principal axes of reconstructed / true point clouds; bottom schematics show deviation from the fronto-parallel reference (slant 0^◦^). (C) Whole-VC slant prediction for each bar type—thin bar, thick bar, cylinder, and thin bar (vertical)—plotted as reconstructed slant (PCA-derived principal-axis angle) against stimulus slant. Each dot is one subject at one stimulus slant angle; gray lines are fitted regression lines; asterisks indicate significant positive slopes (^∗^*p <* 0.05; ^∗∗^*p <* 0.01; ^∗∗∗^*p <* 0.001). Whole-VC slope estimates are reported in Tables S1 (frequentist) and S2 (Bayesian).

We tested slant reconstruction with horizontally elongated bars rotated around the vertical axis and vertically elongated bars rotated around the horizontal axis, each at nominal slants from ±15^◦^ to ±60^◦^ in 15^◦^ steps (these nominal angles map to somewhat shallower effective stimulus slants—e.g., a nominal 60^◦^ corresponds to roughly 50^◦^—which we use as *θ*_stim_ and plot in Figure 5C; see Methods). Because the perceptual reliability of the intended slant varied across subjects at the largest nominal slant (Methods, Perceptual verification), we did not regard the choice between including and excluding it as clear-cut and conducted the slant-prediction analysis both ways: a main analysis excluding the largest-slant condition, reported here, and a parallel sensitivity analysis including it (Figure S7E). The two versions yielded closely matching slope estimates, indicating that the conclusions are robust to this choice.

To obtain a global slant measure insensitive to local surface details, we applied principal component analysis (PCA) to each reconstructed point cloud and took the orientation of the first principal component as the reconstructed structure’s principal axis. Example whole-VC reconstructions and their principal axes are shown in Figure 5B, with the remaining subjects in Figure S5C. The reconstructed principal axis captured the objects’ slant in 3D space for both bar orientations and was visually well aligned with that of the stimulus across slant conditions.

Reconstructed slant broadly followed stimulus slant for every bar type and orientation.

To quantify slant reconstruction, we defined the reconstructed slant as the angle of the principal axis in the appropriate projection of each reconstructed structure and fitted regression lines between stimulus and reconstructed slants. For the thin horizontal bar—our primary slant stimulus, analyzed from the first run type (see Methods)—reconstructions based on the whole VC showed a clear linear relationship between stimulus and reconstructed slants when pooled across subjects, with a reliably positive slope (*β* = 0.77, 95% CI [0.28, 1.27], one-sided *p* = 0.006; Bayesian Pr(*β >* 0) = 0.990; Tables S1–S2). The regression intercept was close to zero, with a slight negative bias (left side of the reconstructed structure closer to the observer). Reliably positive slopes, close to unity, were also obtained for the thick bar (*β* = 1.11, 95% CI [0.64, 1.58], one-sided *p* = 0.001) and the cylinder (*β* = 1.15, 95% CI [0.39, 1.92], one-sided *p* = 0.007; Figure 5C; Tables S1–S2), so slant prediction was robust to bar type. The same conclusions held with the diffusion-based autoencoder (Figure S7F).

These bar types were all horizontally elongated and were acquired in the first run type. To ask whether slant is reconstructed for a different bar orientation as well, we tested a vertical thin bar from a second run type in which horizontal and vertical thin bars were presented together (the horizontal thin bar serving as a matched positive control; see Methods) (Figure 5C). The vertical-bar slope was likewise reliably positive (*β* = 0.37, 95% CI [0.06, 0.68], one-sided *p* = 0.015; Bayesian Pr(*β >* 0) = 0.981; Tables S1–S2), with a near-zero intercept (slight positive bias; lower part of the bar closer to the observer). The same held with the diffusion-based autoencoder (Figure S7F).

Both orientations thus yielded reliably positive slopes. Within the matched runs of the second run type, where both orientations were presented together, the vertical slope tended to be shallower than the horizontal one; we tested this difference as a post-hoc, exploratory analysis (see Methods), but it was neither statistically robust nor qualitatively clear (Δ*β* = 0.16, 95% CI [−0.08, 0.40], *p* = 0.090; Bayesian Pr(*β_H_ > β_V_*) = 0.904), and we do not interpret it further.

These findings show that the pipeline predicts slant from brain activity even when the 2D contour is held constant across stimuli, confirming that 3D reconstruction does not rely on 2D pictorial cues.

### Regional contributions across visual cortex

We evaluated reconstructions from four visual regions—the early VC, MT & neighbors, dorsal VC, and ventral VC (Figure 6A)—to ask how each of the three reconstruction-level tests above (reconstruction of natural objects across trained and novel categories, cross-cue generalization to RDSs, and slant prediction from contour-matched RDSs) depends on cortical region. The regional results are shown in Figures 6 and 7, with the corresponding LMM statistics reported in Tables S3 (identification: per-ROI simple effects and the ROI × stimulus interaction relative to early VC) and S4 (slant slope: per-ROI simple slopes and the ROI × *θ*_stim_ interaction relative to early VC), given as frequentist estimates with Bayesian Pr(dir.); the per-ROI accuracies and slant slopes themselves are shown graphically in Figures 6D–E and 7B.

**Figure 6.**
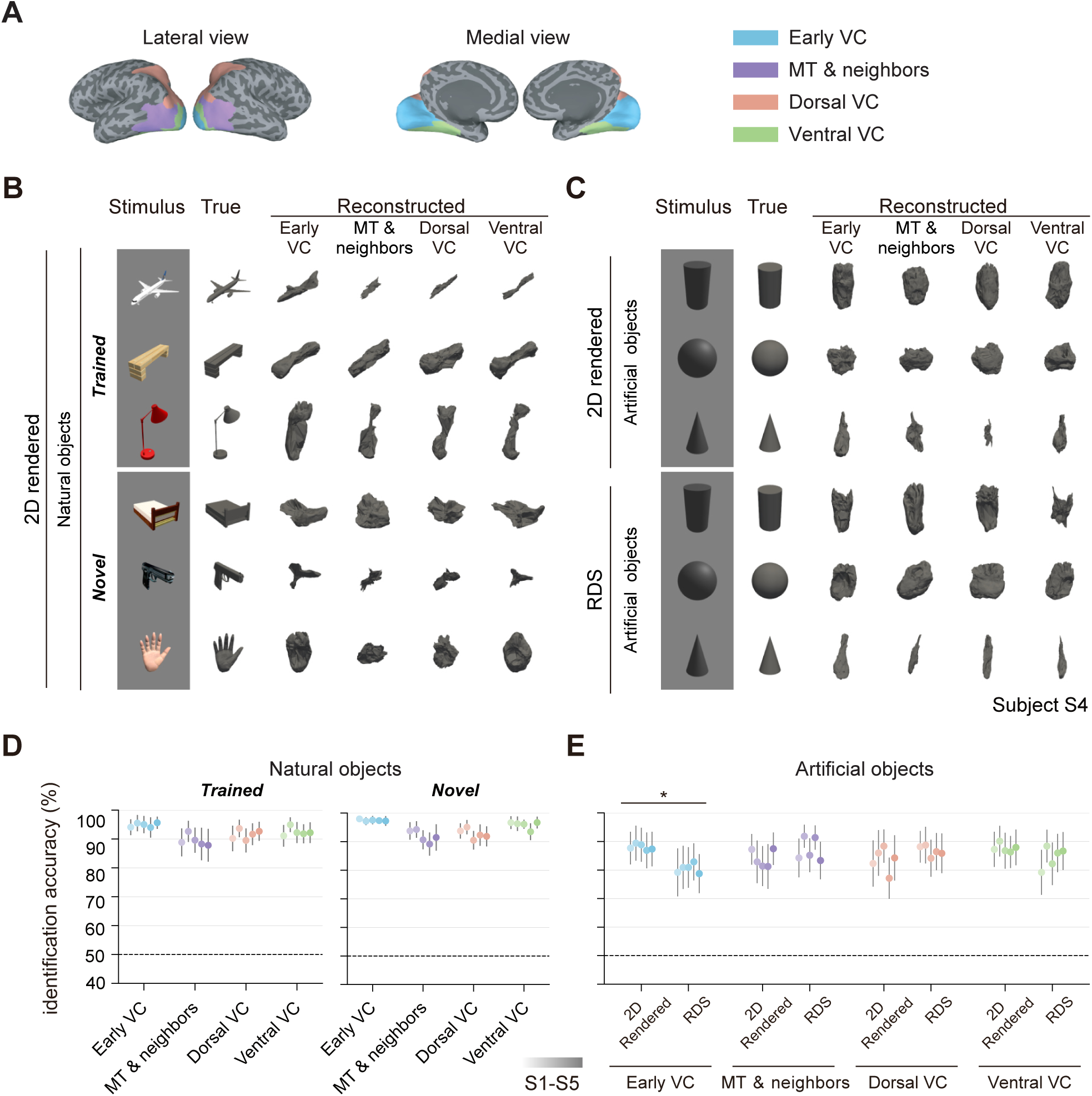
Regional contributions to reconstruction and identification. (A) Four visual-cortex ROIs on the cortical surface. (B) Per-ROI example reconstructions of natural objects (2D rendered images) for subject S4, grouped by category; each row shows the stimulus, ground-truth, and per-ROI reconstructed 3D structures. (C) Same format as (B) for artificial objects under 2D rendered images and RDSs. (D) Identification accuracy across ROIs for natural objects (trained vs novel categories). (E) Identification accuracy across ROIs for artificial objects under 2D rendered vs. RDS conditions; asterisks mark within-ROI differences between the 2D rendered and RDS conditions (per-ROI simple effects from the joint 4-ROI LMM; ^∗^*p <* 0.05); the regional difference is formally assessed by the ROI × stimulus interaction (Table S3). In (D) and (E), each dot is one subject (S1–S5); error bars are 95% CIs; dashed line indicates chance (50%).

**Figure 7.**
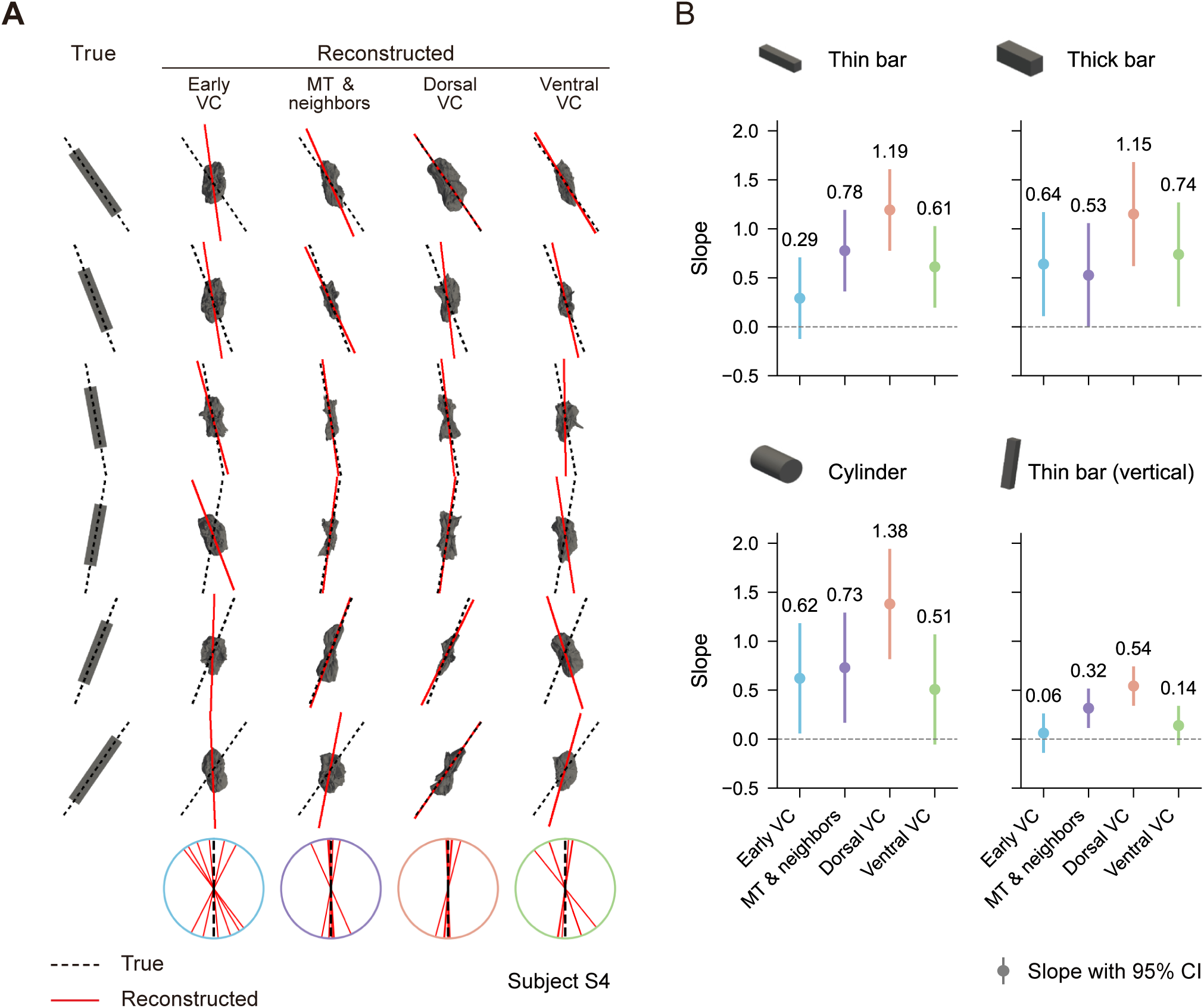
Cross-ROI slant prediction from contour-matched RDSs. (A) Per-ROI example reconstructions of a horizontal bar (contour-matched RDS) for subject S4 across stimulus slants. Red solid / black dashed lines: principal axes of reconstructed / true point clouds; bottom schematics show principal-axis deviation from the fronto-parallel reference (as in Figure 5B). (B) Forest plots of ROI-wise slant-prediction slope for each bar type—thin bar, thick bar, cylinder, and thin bar (vertical); points are per-ROI simple slopes from the joint 4-ROI LMM and horizontal lines are 95% CIs; the regional difference is formally assessed by the ROI × slant interaction. Per-ROI simple slopes (one-sided test of slope *>* 0) and the interaction relative to early VC are reported in Table S4.

For 2D rendered natural objects, all ROIs achieved high identification accuracy (*>* 80%), with the early VC highest, the dorsal and ventral VCs comparable, and the MT & neighbors lowest (Figure 6D; above chance in every subject). A similar regional pattern was observed with the diffusion-based autoencoder (Figure S4C).

Identification accuracy for RDSs was above chance in every region and every subject (Figure 6E). Examined region by region, identification accuracy was comparable across the two stimulus formats in the higher-order ROIs (MT & neighbors, dorsal VC, and ventral VC), whereas in the early VC it was lower for RDSs than for 2D rendered images (Figure 6E; per-ROI simple effect in Table S3). The regional difference was established by a ROI × stimulus interaction analysis: the drop from 2D rendered images to RDSs was significantly larger in the early VC than in MT & neighbors and dorsal VC (MT & neighbors vs. early VC: *β*_interaction_ = +10.7%, 95% CI [+4.8, +16.5], *p <* 0.001; dorsal VC vs. early VC: *β*_interaction_ = +10.6%, 95% CI [+4.7, +16.4], *p <* 0.001; Table S3). Under the diffusion-based autoencoder, identification accuracy showed the same qualitative pattern, with the early VC again lower for RDSs (Figure S4D).

For contour-matched RDSs (Figure 7), slant prediction was strongest in the dorsal VC, followed by MT & neighbors and the ventral VC, and weakest in the early VC. Within each region, the per-ROI slant slopes (simple slopes; Figure 7B) showed that the dorsal VC had reliably positive slopes for every bar type—thin, thick, cylinder, and vertical (all *p <* 0.001)—making it the most robust region for slant prediction, whereas the early VC reached significance for neither the horizontal nor the vertical thin bar. The difference between regions was then assessed by the ROI × slant interaction relative to early VC, the same form of test as for identification (Table S4): the slant slope was significantly steeper in the dorsal VC than in the early VC for the horizontal thin bar, the vertical thin bar, and the cylinder (Δ*β* = +0.90, 95% CI [+0.54, +1.26], *p <* 0.001; Δ*β* = +0.48, [+0.27, +0.69], *p <* 0.001; and Δ*β* = +0.76, [+0.09, +1.43], *p* = 0.026, respectively), with a positive but non-significant difference for the thick bar (Δ*β* = +0.51, *p* = 0.078); MT & neighbors was also significantly steeper than the early VC for the horizontal and vertical thin bars (Δ*β* = +0.49, [+0.13, +0.84], *p* = 0.008, and Δ*β* = +0.25, [+0.05, +0.46], *p* = 0.017), but not for the thick bar or cylinder (both *p* ≥ 0.69). This thin-bar specificity reflects the early VC’s own slant tuning: its slope was near zero for the thin bars but non-negligible and significant for the thick bar and cylinder. All bar types were equally contour-matched, so binocular disparity was the only slant cue throughout; the thick bar and cylinder nonetheless differ from the thin bar in more than size, carrying additional disparity-defined structure such as surface curvature and extended edges, and we therefore do not draw strong mechanistic conclusions from the differences among bar types. The robust observation is that the advantage of the dorsal VC and MT & neighbors over the early VC emerged most clearly for the thin bar. Crucially, the dorsal VC gave the steepest slope for every bar type (Figure 7B), so its advantage did not depend on the particular shape or orientation of the bar; the relative ordering of the early VC, MT & neighbors, and ventral VC, by contrast, varied across bar types. The exploratory comparison of horizontal versus vertical thin-bar slopes showed the same direction in every ROI (slope tended to be shallower for vertical), but a reliable difference appeared only in MT & neighbors (Δ*β* = 0.26, one-sided *p* = 0.013; Bayesian Pr(*β_H_ > β_V_*) = 0.986; Table S4); the other ROIs (all *p* ≥ 0.10; Table S4) and the whole VC (one-sided *p* = 0.090; Tables S1 and S2) showed no reliable difference. As an exploratory effect, we do not interpret it further.

Together, these results reveal a regional dissociation. From 2D rendered images, the identification accuracy of the resulting 3D structures was high across all ROIs, tending to be strongest in the early and ventral VCs; this identification accuracy generalized to RDSs in all regions except the early VC, which tended to drop. The regional differences were far more pronounced for slant prediction. Because contour-matched RDSs hold the 2D outline fixed and define slant by binocular disparity alone, accurate slant prediction—where training and test necessarily differ in depth cue—provides a direct test of a cue-invariant 3D representation. Slant was predicted most strongly and consistently in the dorsal VC (significant for every bar type), with MT & neighbors also reliable, whereas the early VC was weakest: it failed specifically for the thin bars while still showing slant sensitivity for the thick bar and cylinder. The dorsal VC and MT & neighbors thus most clearly carry the cue-invariant 3D representation, with the early VC contributing least, particularly for the thin bars.

## Discussion

We show that an internal, cue-invariant representation of 3D structure can be externalized from human brain activity as an explicit 3D object. Latent features of a 3D point-cloud autoencoder were reliably decoded from visual cortical responses to 2D rendered images and used to reconstruct 3D structures that preserved coarse shape and 3D orientation. The same decoder— trained exclusively on fMRI responses to 2D rendered images—generalized to brain activity elicited by random dot stereograms (RDSs) that contain no pictorial depth cues, and further reconstructed 3D structures from contour-matched RDSs, in which the 2D contour was held constant and only the 3D structures varied. Cross-cue generalization was more pronounced in higher visual areas than in early visual cortex.

These results show that cue-invariant 3D structure can be externalized from human fMRI responses as a directly inspectable 3D object. Previous similarity-based analyses of cortical response patterns could test whether representations are shared across cues, but did not make the 3D structure that the brain constructs from those cues directly inspectable.

The cross-cue results further clarify the structure of cortical 3D representations. A substantial portion of reconstruction performance transferred from rendered-image training to RDS test stimuli, indicating a cue-invariant component that abstracts away from the specific retinal cue conveying depth. The clearest example is slant reconstruction from contour-matched RDSs, whose 2D outlines are identical across slants and share no pictorial shape information with the rendered-image training set, so that successful slant reconstruction cannot be attributed to 2D contour information. Cross-cue generalization therefore most directly exposes this cue-invariant component of cortical 3D representations.

The regional pattern points most clearly to the dorsal VC as a contributor to this cue-invariant component, with MT & neighbors also implicated (Figures 6 and 7). For identification, the drop in accuracy from 2D rendered images to RDSs was significantly smaller in both the dorsal VC and MT & neighbors than in the early VC, whereas for slant the advantage over the early VC was clearest and most consistent in the dorsal VC, which was significantly steeper than the early VC for the thin bars and the cylinder; MT & neighbors was also significantly steeper than the early VC for the thin bars. Rendered images and RDSs also differ in low-level properties unrelated to depth cues (see Methods, Visual stimuli), which may partly account for the weaker generalization in early VC. However, such differences cannot explain the successful generalization in higher visual areas, making a purely low-level account unlikely. This regional dissociation is broadly consistent with previous electrophysiological and neuroimaging work: the early visual cortex (V1, V2) processes individual depth cues such as disparity and texture without clear evidence of cue-invariant, perceptually relevant 3D coding^25–28^, whereas higher dorsal regions (overlapping with V3A and V3B/KO in our dorsal VC) and ventral object-selective regions support disparity-based and cue-integrated 3D representations^29–32^. Our results extend these findings by showing that representations in these higher visual areas can support reconstruction of explicit 3D shape, not merely categorical or relational depth information.

The slope relating reconstructed to stimulus slant varied with bar type: it was close to unity for the thick bar and cylinder and positive though shallower for the thin bars, especially the vertical thin bar (Tables S1 and S2). Several factors could shape these slope values. The decoding and generation pipeline is itself noisy and constrained—voxel patterns are decoded into latent features, a nonlinear generator maps these features to a point cloud, and PCA then reads out the slant—and perceptual factors, such as well-documented individual variation and biases in perceived surface slant^33–36^, may also contribute; our design does not separate these contributions. In a matched comparison, the horizontal–vertical slope difference was modest and only exploratory (Results), so we draw no firm conclusion from it; the tendency toward shallower reconstructed slant for vertical bars is nonetheless in the direction expected if anisotropic perceptual priors—shaped by the differing slant statistics of horizontal and vertical surfaces in natural scenes^37–39^—bias the reconstructed surface slant.

Several limitations define the scope of these conclusions. Because the study used a small number of intensively sampled subjects and a constrained object/stimulus set, our claim concerns reproducible within-subject reconstruction of controlled 3D geometry, not a general-purpose decoder of arbitrary 3D percepts. Three limitations are especially important.

First, the experimental design and pipeline restrict what can be reconstructed. The stimuli were single isolated objects rendered against a blank surround, and the pipeline therefore reconstructs only 3D geometry devoid of material or photometric attributes. The 3D autoencoders we used operate on point clouds without surface texture or reflectance; extending the pipeline to generate realistic textures, translucency, and view-dependent appearance will require generators that jointly model structure and surface properties and/or flexible 3D representations such as NeRF^40^ and Gaussian splatting^41^. The generalization from 2D rendered images to RDS stimuli also involved low-level differences beyond depth-cue type (see Methods, Visual stimuli), so the weaker reconstruction performance to RDSs observed in the early visual cortex does not immediately rule out cue-invariant representations in this region. A further scope constraint lies in the architecture of the 3D output space. AtlasNet—our primary backbone—generates surfaces by deforming a fixed template (a single sphere in our configuration) through a learned mapping conditioned on a latent code^22^. The representable outputs are therefore confined to closed, genus-zero surfaces obtained from one primitive, and cannot capture multi-object scenes, occlusion, surfaces with holes or branching topology, or disconnected parts. This is a substantially narrower prediction space than the pixel-level outputs targeted by 2D image-reconstruction pipelines. The training regime has the same scope: both the 3D autoencoder and the brain-feature decoder were trained on single-object stimuli on a blank surround, with no exposure to multi-object compositions or natural scene structure. Extensions to multi-object scenes are therefore a natural direction for future work and are discussed below.

Second, the statistical scope is limited by the sample of five subjects. This sample size limits the number of grouping levels for the random effects in the linear mixed-effects models, which may reduce statistical power for detecting smaller effects and lead to imprecise estimates of between-subject variance. It reflects an intensive within-subject design—widely advocated in cognitive neuroscience and central to our prior work—in which each subject is treated as an independent replication of the experiment^42,43^; the small subject count is therefore a deliberate consequence of dense per-subject sampling rather than an under-powered group study. Reassuringly, the central results were quantitatively consistent across all five subjects: per-subject identification accuracies (Figures 3F, 4F, 6D, 6E), Chamfer distances (Figure S6B), and slant-prediction patterns (Figure S5C) showed broadly similar cross-subject patterns. We further addressed the small-grouping-level limitation by applying the Kenward–Roger small-sample correction in the main frequentist analysis, and by running a parallel Bayesian analysis with weakly informative priors as a robustness check; the qualitative pattern of results was consistent across the two frameworks.

Third, the success of the pipeline should not be taken as evidence that cortical 3D representations match the specific latent features used here. The present study examined latent features from two point-cloud autoencoders that share the same input format, encoder architecture, and training dataset; they are likely to yield similar latent feature representations, and the results do not imply that 3D representations in the brain are particularly similar to these specific features.

More fundamentally, our reconstructions may partially reflect memorization and interpolation within the training shape manifold rather than truly compositional 3D reconstruction from primitive bases. Although we evaluated category generalization (Figures 3 and 4), the training set covers a broad fraction of common everyday object geometries, so a system that internalizes the training shape manifold and retrieves a nearby exemplar followed by mild interpolation could plausibly account for part of the observed performance. Two recent analyses of brain-to-stimulus reconstruction pipelines crystallize this concern: Otsuka et al.^44^ show that linear translators in brain-decoding pipelines can exhibit *output dimension collapse*, in which decoded outputs are confined to the low-dimensional subspace spanned by training-feature clusters, and Shirakawa et al.^20^ argue that apparent reconstruction success can arise from category-level classification followed by generative completion rather than from faithful reconstruction of the specific content. Whether our pipeline performs genuinely *compositional* prediction—reconstructing arbitrary 3D shapes by composing learned 3D primitives or bases, rather than retrieving and interpolating over a single-object training manifold—therefore remains uncertain. Resolving this question, together with the experimental-scope limitations identified above, points to extending the translator-generator framework to multi-object scenes as a particularly informative direction for future work. Informative tests for compositionality more specifically include reconstructions of objects with non-sphere topology and of stimuli placed explicitly outside the convex hull of the training feature distribution.

Placed in the broader trajectory of brain decoding, the present study extends reconstruction from retinal-format outputs to explicit 3D structure. Brain decoding has progressed from reading out the orientation of a viewed grating from primary visual cortex activity^45^ to DNN-based reconstruction of perceptual experiences^13–15,46^ (see^47^ for a recent review). Most of this work has focused on retinal-format outputs—2D images or spectrograms. Initial steps toward explicit 3D output have been reported^19^, but, as discussed in the Introduction, these efforts have relied heavily on 2D image and semantic features and have not tested whether the reconstruction reflects a cue-invariant 3D representation. Our results extend the cross-decoding approach to explicit 3D structure and add the missing cue-invariance test: the latent features of pretrained 3D point-cloud autoencoders are aligned with cortical 3D representations well enough to support reconstruction of coarse 3D structure across the tested depth cues.

This extension also links reconstruction to the idea of an internal world model. Unlike 2D image reconstructions, which reproduce the momentary retinal appearance, an explicit 3D reconstruction externalizes the structure that the brain would use to predict the sensory consequences of viewpoint changes and interactions. In this sense, decoding explicit 3D structure from brain activity can be regarded as a readout of the brain’s internal world model—the internal representation that the brain infers from sensory inputs and uses to anticipate how the same content would appear under changes of viewpoint or other actions, and that is intrinsically 3D rather than retinal^11,12^. In our study, this readout generalized across qualitatively different depth cues, indicating that the framework accessed visual-cortical information consistent with a common 3D representation, a component of what may support an internal world model. The present study demonstrated such integration within the visual modality, but the brain also combines inputs across vision, touch, audition, and other modalities to build a coherent world model. Whether our approach can externalize such a multimodal world model, and which DNN features best capture the brain’s 3D representations—across architectures, training objectives, and coordinate systems (e.g., viewer-centered vs. object-centered)—are important open questions for future research.

In conclusion, cue-invariant internal 3D representations can be externalized from human brain activity in an explicit format that makes internal 3D structure directly inspectable. This bridge between neural activity and reconstructed 3D structure provides a route for probing how the brain constructs 3D object representations and for asking how it assembles a broader internal model of the world.

## Resource availability

### Lead contact

Further information and requests for resources should be directed to and will be fulfilled by the lead contact, Yukiyasu Kamitani.

### Materials availability

This study did not generate new unique reagents.

### Data and code availability

- All data reported in this paper are shared at figshare (https://doi.org/10.6084/m9.fig share.c.8508462).
- Raw fMRI data are shared at OpenNeuro (https://openneuro.org/datasets/ds007792).
- Code to reproduce the main results is shared at GitHub (https://github.com/KamitaniLab/3dReconstruction).

## Acknowledgments

This study was supported by Japan Society for the Promotion of Science (KAKENHI grants JP20H05705, JP25K24743, and JP20H05954 to Y.K.), the New Energy and Industrial Technology Development Organization (grant JPNP20006 to Y.K.), and Japan Science and Technology Agency (CREST grant JPMJCR22P3). The MRI experiments were conducted at the Kyoto University Institute for the Future of Human Society with technical support from the institute’s staff. We are deeply grateful to Mitsuaki Tsukamoto for his major contributions to the inception and early development of this project. We thank Kenya Otsuka and Taiga Kurosawa for their assistance with the experiments.

## Use of generative AI

During the preparation of this work, the authors used Claude, ChatGPT, and Gemini to revise manuscripts and edit code. After using these tools, the authors reviewed and edited the content as needed and took full responsibility for the content of the publication.

## Supplementary information

Figures S1–S7 and Tables S1–S4 with their legends are provided in the Supplementary Information section at the end of this document.

## Methods

### Subjects

The dataset presented in this paper is from five subjects with normal or corrected-to-normal vision (four males and one female; aged 23 to 40 years). All subjects provided informed consent prior to the experiment. The study protocol and the instruments used in MRI experiments were approved by Kyoto University (approval no. KUIS-EAR-2017-002) and the Ethics Committee of the Advanced Telecommunications Research Institute International (approval no. 106), and was conducted following the principles of the Declaration of Helsinki.

### Visual stimuli

We presented the subjects with two classes of visual stimuli: (1) 2D rendered images of 3D objects and (2) random dot stereograms (RDSs) that elicit 3D perception via binocular disparity.

### 2D images

2D images were created from two domains: 1) natural objects and 2) artificial objects. All 2D images were rendered with Blender as 1024 × 1024-pixel RGBA images.

#### Natural objects

We collected 3D meshes of everyday objects from ShapeNet^48^ and SketchUp 3D Warehouse (https://3dwarehouse.sketchup.com/). As training stimuli, we selected 1,900 objects spanning 19 categories (100 objects per category) from ShapeNet and 100 objects of animals from 3D Warehouse, resulting in a set of 2,000 objects across 20 categories (“airplanes”, “benches”, “cabinets”, “cars”, “chairs”, “displays”, “lights”, “speakers”, “rifles”, “sofas”, “household containers”, “bathroom fixtures”, “bowls and jars”, “buses and trains”, “electronic and digital devices”, “home appliances”, “guitars”, “tables”, “watercrafts”, and “animals”). Categories from ShapeNet were either 1) the original ShapeNet label or 2) merged ShapeNet labels that were either semantically or geometrically similar (e.g., “buses” and “trains”). As test stimuli, we additionally selected 72 objects spanning 18 categories (4 objects per category) from ShapeNet and 8 objects of “hands” or “human figures” (4 objects for each) from 3D Warehouse, resulting in 80 objects across 20 categories. Ten categories (“airplanes”, “benches”, “cabinets”, “cars”, “chairs”, “displays”, “lights”, “speakers”, “rifles”, and “sofas”) were shared between the training and test stimuli, but individual objects did not overlap. The remaining ten categories (“beds”, “bottles”, “caps”, “helmets”, “knives”, “motorbikes”, “pianos”, “pistols”, “hands”, and “human figures”) were included only in test stimuli. Because individual objects never overlapped between the training and test sets, every test object was a new exemplar; the two groups of natural test objects therefore differ only in whether the object’s category was present during training. The 40 test natural stimuli from the shared categories were referred to as “trained categories”, and the remaining 40 from categories absent from the training set as “novel categories”.

For the training set, objects were rotated uniformly at random in the 3D space. For the test set, objects were rotated only around the vertical axis and rendered as they were viewed from a slightly elevated top-down perspective. The viewpoints were adjusted to exclude accidental views or abnormal appearance in which the 3D structure was ambiguous to observers. All natural objects were rendered with surface color and texture included in the 3D mesh data.

#### Artificial objects

As abstract 3D objects without semantic categorization, we created 16 geometric primitives (cube, bar, sphere, ellipsoid, cylinder, bent cylinder, cone, truncated cone, pyramid, octahedron, torus, hourglass, disk, square plate, triangular plate, and L-shaped tetracube) using Blender. We rotated some primitives to produce view variants in which the same primitives appeared as different 3D structures from the viewer’s perspective (2 views for cone, pyramid, disk, square plate, and triangular plate; 3 views for bar, cylinder, and hourglass; 4 views for bent cylinder). We treated the differently rotated primitives as distinct 3D structures in this study. Thus, the artificial objects consisted of 30 stimuli in total. The surface color was dark gray for all artificial objects. All artificial objects were rendered from a slightly elevated top-down perspective.

### Random dot stereogram

RDSs were created for two domains, artificial objects and slant bars. For each 3D object, we obtained a depth map in Blender and converted it to a binocular disparity field for a virtual observer with a 60 mm inter-ocular distance, interpreting Blender coordinates as meters. The reference (zero-disparity) viewing distance for this conversion was set per object from its depth in the scene (a few meters) and therefore differed from the physical presentation distance (≈ 0.82 m; see Contour-matched RDSs for the consequences); disparity was zero at the center of the field, where subjects fixated. The field was applied to a pair of random dot patterns (50% white, 50% black) by shifting dots (dot size 0.15^◦^, density 75%); the full construction is in the code repository (see Resource availability). The background was a fixed-disparity RDS (uncrossed disparity 0.57^◦^, ≈ 3 m in the Blender scene) for all stimuli.

#### Artificial objects

We used the same 30 artificial objects used for 2D images for RDSs.

#### Contour-matched RDSs

These stimuli were designed so that the cyclopean (single, central-view) silhouette—the projection of the 3D structure onto the fronto-parallel plane—was identical across stimuli, while the disparity-defined slant differed. Matching is on this cyclopean outline, not on the two eyes’ retinal images: the left- and right-eye half-images still differ, and that interocular disparity is what conveys slant. Because the outline (and hence each bar’s projected thickness) is held constant across slants, the stimuli carry no contour- or size-based pictorial depth cue, so binocular disparity is the only cue to slant. We used three primitives—thin bar, thick bar, and cylinder—with thick bars and cylinders lying horizontally and thin bars oriented either horizontally or vertically. For a horizontal bar, depth shifts the horizontal offset of the bar’s ends between the two eyes; for a vertical bar, it changes the bar’s interocular tilt; in both cases the projected thickness stays uninformative about depth. Each primitive was slanted by nominal angles of ±15^◦^, ±30^◦^, ±45^◦^, and ±60^◦^ (horizontal bars around the vertical axis, vertical bars around the horizontal axis) in Blender and then manually deformed until its cyclopean silhouette matched pixel-for-pixel across slants; the resulting point clouds are deposited on figshare (see Resource availability). These angles are the Blender rotation angles set at design time. Because the screen size and viewing distance assumed when generating the disparities were imprecise and varied across stimuli (presentation distance ≈ 0.8 m versus a per-object generation distance of a few meters), the slants actually specified by the on-screen disparity were somewhat shallower than these nominal values and varied across stimuli: the rescaling acts as a multiplicative gain of roughly 0.6–0.7 on the tangent of the slant (tan *θ*_actual_ ≈ 0.6–0.7 × tan *θ*_nominal_), so on average the nominal ±60^◦^, ±45^◦^, ±30^◦^, and ±15^◦^ bars corresponded to effective slants of roughly ±50^◦^, ±35^◦^, ±20^◦^, and ±10^◦^, respectively. We therefore recomputed each stimulus’s actual geometric slant from the presentation geometry and used it as the stimulus slant (*θ*_stim_) in all analyses (nominal angles gave essentially the same results). This *θ*_stim_ is a geometric quantity and need not equal the perceived slant, which may vary across observers; because the 2D contour is held constant, none of this affects the contour-matching logic.

### fMRI experiments

#### Experimental design

We conducted visual stimulus presentation experiments for two types of fMRI sessions: training and test sessions. Visual stimuli were rear projected onto a screen inside the bore of an MRI scanner with a gamma-corrected projector (DLA-V70R, Victor, Yokohama, Japan). The visual stimuli were presented with Matlab (version 2023a) and Psychtoolbox (version 3.0.19). In RDS experiments, the stimuli were presented as anaglyphs using stereoscopic stimulus presentation implemented in Psychtoolbox. During the RDS experiments, subjects wore red-cyan anaglyph glasses. The stimuli were displayed at the screen center with a size of 12^◦^ × 12^◦^ of visual angle on a gray, mid-luminance background (or, in RDS experiments, on a fixed-disparity random-dot stereogram background; see Visual stimuli). We asked subjects to fixate on the center of the screen cued by a circle during the experiment. To reduce head motion during fMRI data collection, subject S1 used a personalized headcase (https://github.com/gallantlab/head case-pipeline), and subjects S2–S5 used a custom-molded bite bar.

#### Training session

A single set of training sessions consisted of presenting each of the 2,000 training stimuli once in 40 runs. Each run contained 55 trials, with 50 trials of different stimuli and five randomly interspersed one-back repetition trials that showed the same image as the previous trial. In each trial, a stimulus was flashed at 1 Hz for 8 s. At the beginning and end of each run, 32-s and 6-s rest periods were added, respectively. During the rest period, only the fixation point was presented. There was no inter-trial interval. We instructed the subjects to maintain fixation at the fixation point throughout the run and to perform a one-back repetition detection task; the subjects were required to press a button with their right hand when the identical stimulus was repeatedly presented in successive trials. The data from the repetition trials were excluded from the analyses. We repeated the single set of training sessions three times for each subject, resulting in a total of 120 runs.

#### Test session

In the test sessions, each test stimulus set (2D rendered images of natural objects, 2D rendered images of artificial objects, RDSs of artificial objects, and RDSs of slant bars) was presented in separate runs. The time course of the test runs (trial duration and rest periods) was identical to that of the training sessions. 2D rendered images were presented flashed at 1 Hz. RDSs were presented continuously as static random dot stereograms for the entire trial duration (no temporal flashing). Thus, 2D rendered images and RDSs differed in several low-level properties beyond depth-cue type: temporal structure (1 Hz flashing vs. continuous presentation), chromatic format (RGB image vs. red-cyan anaglyph image), and background composition (uniform gray vs. random dot pattern). These differences are considered when interpreting the generalization results (see Discussion). Subjects were instructed to fixate on the central fixation point and to perform the same one-back repetition detection task as in the training sessions. Randomly interspersed one-back repetition trials were included within each run, and the data from these trials were excluded from the analyses.

For the 2D images of natural objects, 80 stimuli were presented in two runs and repeated eight times. Each run included five one-back repetition trials. A total of 16 runs were collected across two scanning sessions.

For the 2D images of artificial objects, 30 stimuli were presented in a single run and repeated eight times. Each run included three one-back repetition trials. A total of eight runs were collected across one scanning session.

For RDSs of artificial objects, 30 RDS stimuli were presented together with five 2D rendered images of the test natural objects. These 2D rendered images were included so that the per-voxel mean response across the whole run—which served as the baseline for the response estimates—was stabilized and kept comparable to that of the other runs; they were presented on the same random-dot background as the RDS stimuli and were not used in the analyses. A total of 35 stimuli were presented in a single run and repeated eight times. Each run included four one-back repetition trials. A total of eight runs were collected across one or two sessions.

For the RDSs of slant bars, two types of runs were used. In the first type, three bar types (thin bars lying horizontally, thick bars, and cylinders) were presented with eight nominal slant angles (±60^◦^, ±45^◦^, ±30^◦^, and ±15^◦^ nominal), resulting in 24 RDS stimuli. Additionally, 24 2D rendered images of test natural and artificial bar stimuli were included to stabilize the run-wise baseline (the per-voxel mean response across the run, as above) and were presented on the same random-dot background as the RDS stimuli, yielding a total of 48 stimuli presented in a single run; these images were not used in the analyses. Each run included five one-back repetition trials. In the second type, only thin bars were used, presented in two orientations (horizontal and vertical) and slanted with the same eight angles, resulting in 16 RDS stimuli. Additionally, 16 2D rendered images of test stimuli were included to stabilize the run-wise baseline (as above) and were presented on the same random-dot background as the RDS stimuli, yielding a total of 32 stimuli presented in a single run; these images were not used in the analyses. Each run included four one-back repetition trials. Each run was repeated eight times. A total of 16 runs were collected across two sessions. For the slant analysis, the fMRI responses to each stimulus were averaged across its eight repetitions. Per-bar-type slope estimates were computed from the first run type for the horizontal thin bar, thick bar, and cylinder, and from the second run type for the vertical thin bar. For the direct comparison between the horizontal and vertical orientations, both thin bars were instead taken from the second run type, in which the two orientations were presented together under matched conditions; the horizontal thin bars of the second run type were otherwise treated as controls and did not enter the per-bar-type estimates.

#### Perceptual verification

After all experimental sessions were completed, we conducted a debriefing in which subjects verbally reported whether they had perceived the intended 3D structure for each stimulus type. For the 2D rendered images and the artificial-object RDSs, all five subjects confirmed that they had perceived clear 3D structure. For the contour-matched RDSs, two of the five subjects reported that at the largest nominal slant (±60^◦^ nominal), the bar appeared distorted or that they could not reliably perceive the intended slant. Because we could not determine on the basis of these reports alone whether excluding these conditions was clearly warranted, we ran the slant-prediction analysis under both choices—on the subset excluding them (reported as the main analysis) and on the full set including them (reported as a sensitivity analysis; Figure S7E)—and confirmed that the two analyses yielded closely matching slope estimates, indicating that the conclusions are robust to this choice.

### MRI acquisition

We collected fMRI data using a 3.0-Tesla Siemens MAGNETOM Verio scanner at the Kyoto University Institute for the Future of Human Society. An interleaved T2*-weighted gradient-echo echo planar imaging (EPI) sequence was used to acquire whole-brain functional images (TR 2000 ms, TE 43 ms, flip angle 80^◦^, FOV 192 × 192 mm^2^, voxel size 2 × 2 × 2 mm^3^, slice gap 0 mm, number of slices 76, multiband factor 4). At the beginning of each scanning session, prior to the functional runs, we acquired a field map consisting of two magnitude images and one phase difference image using a double-echo gradient-echo sequence for distortion correction of the functional EPI images (long TE 12.46 ms, short TE 10 ms, TR 1200 ms, flip angle 90^◦^, FOV 192 × 192 mm^2^, voxel size 2 × 2 × 2 mm^3^, number of slices 76). High-resolution T1-weighted anatomical images of the entire head were acquired from each subject prior to the main experiment using a magnetization-prepared rapid acquisition gradient-echo (MP-RAGE) sequence (TR 2250 ms, TE 3.06 ms, TI 900 ms, flip angle 9^◦^, FOV 256 × 256 mm^2^, voxel size 1.0 × 1.0 × 1.0 mm^3^, number of slices 208).

### fMRI preprocessing

We preprocessed the MRI data through an in-house pipeline based on fMRIPrep^49^ (version 1.2.1). Before the preprocessing of the functional data, we created an anatomical reference T1w image for each subject from the MP-RAGE fine-structural T1w image (1 mm isotropic voxels). Bias correction using SPM (version 12) was applied to the T1w image. For the functional data of each run, we first estimated a BOLD reference image using the standard methodology implemented in fMRIPrep. Then, susceptibility distortion correction with field maps was performed using FSL. The data were subsequently motion-corrected using MCFLIRT from FSL (version 5.0.9) and slice time-corrected using 3dTshift from AFNI (version 16.2.07), based on the BOLD reference image. Next, we coregistered the anatomical reference T1w image using boundary-based registration implemented by bbregister from FreeSurfer (version 6.0.1). The coregistered BOLD time series were then resampled onto their original space (2 mm isotropic voxels) with antsApplyTransforms from ANTs (version 2.1.0) using Lanczos interpolation. We further preprocessed the fMRIPrep outputs to generate trial-wise response data. First, we shifted the BOLD time series by 4 s (two volumes) to compensate for hemodynamic delays. Then, we regressed out nuisance parameters from each voxel’s time series of each run, including a DC component, a linear trend, and temporal component proportional to the six head-motion parameters estimated during the head motion correction by fMRIPrep. Next, we reduced outliers of the time series (beyond ±3 SD for each run). Finally, we averaged the signal values within each 8-s trial (four volumes) to obtain fMRI response for each stimulus presentation.

### Regions of interest

We defined regions of interest (ROIs) in the visual cortex based on the multi-modal cortical parcellation from the Human Connectome Project (HCP MMP1.0)^21^. To adapt the HCP parcellation onto individual brains, we first mapped HCP MMP1.0 parcellation onto fsaverage cortical surface (https://wiki.humanconnectome.org/docs/assets/Resampling-FreeSurfer-HCP_5_8.pdf), and converted the labels into surface annotation. Then, we reconstructed each subject’s cortical surfaces from their anatomical reference T1w image using FreeSurfer (version 7.2.0). The HCP MMP1.0 annotation was projected from fsaverage to each individual’s cortical surfaces using mri surf2surf from FreeSurfer, and then converted into individual surface labels. These labels were subsequently transformed into volumetric masks aligned to the subject’s anatomical reference T1w image (1 mm isotropic voxel) using mri label2vol from FreeSurfer. The resulting volumetric masks were resampled to the native functional space (2 mm isotropic voxel) using flirt from FSL (version 5.0.2.1).

We defined four visual cortical regions as unions of the HCP parcels: early visual cortex (early VC), MT and its neighbors (MT & neighbors), dorsal visual cortex (dorsal VC), and ventral visual cortex (ventral VC) (Table 1). In addition, we defined “whole visual cortex” (whole VC) as the union of these four regions together with the additional parcels DVT, ProS, POS1, and POS2 (Table 1).

### 3D latent representation

To obtain compact representations of 3D structure suitable as decoding targets, we trained two 3D point-cloud autoencoders that map between explicit 3D structures (point clouds) and low-dimensional latent feature vectors: AtlasNet^22^ and a diffusion-based point-cloud autoencoder^23^. The encoder of each autoencoder yields a latent feature used as the decoding target, and the generator reconstructs a point cloud from a (decoded) latent feature. AtlasNet served as the primary model, and the diffusion-based model was used to confirm that the results do not depend on a particular network framework.

### AtlasNet

We used the autoencoder version of AtlasNet. We used Python implementation of AtlasNet available at https://github.com/ThibaultGROUEIX/AtlasNet for the feature extraction and 3D generation. AtlasNet was trained using the code provided at https://github.com/YefanZhou/d ispersion-score.

The encoder module, based on PointNet^50^, transforms an input point cloud into a 1024-dimensional latent feature. The encoder first maps the input point cloud *x* ∈ R*^N^*^×3^, where *N* is the number of points, onto a 1024-dimensional space via a pointwise multi-layer perceptron, yielding a pointwise feature representation *h* ∈ R*^N^*^×1024^. A channel-wise max pooling across the *N* points then aggregates these pointwise features into a single permutation-invariant global vector *g* ∈ R^1024^. The global vector is further passed through two fully connected layers, each followed by batch normalization and ReLU activation, maintaining the dimensionality at 1024. The final output is the 1024-dimensional embedding *z* ∈ R^1024^ of the input point cloud.

The generator module recovers a 3D structure as a point cloud by deforming one template. The input to the generator module is the 1024-dimensional 3D feature *z* extracted by the encoder module. Given an input vector encoding a 3D structure, the generator, implemented as a multi-layer perceptron, maps the 2D template points to 3D space, conditioned on the latent vector *z*. The output of the generator is a point cloud *p* ∈ R*^M^*^×3^, where *M* is the number of template points. In the current study, we used a single sphere template consisting of 2,562 points.

### Diffusion-based point-cloud autoencoder

The diffusion-based point-cloud autoencoder consists of a PointNet encoder and a diffusion-based decoder. The encoder follows the same architecture as AtlasNet, but produces a 256-dimensional latent feature from an input point cloud.

The generator module recovers a 3D structure as a point cloud using a conditional diffusion probabilistic model, in contrast to AtlasNet’s template-deformation approach. Given the 256-dimensional latent feature, the generator outputs a point cloud through an iterative denoising process of 200 steps with a linear variance schedule (*β* increasing linearly from 1 × 10^−4^ to 0.05). The denoising network is a six-layer MLP. The output of the generator is a point cloud of 2,500 points.

### Model training

We trained point-cloud autoencoders using the ShapeNetCore^48^ v1 dataset, which contains 57,448 3D objects spanning 57 categories. For data augmentation, every object was rotated with a uniformly random 3D rotation. After training, model performance was evaluated on the test objects used in the fMRI experiments.

For AtlasNet, the model was optimized to minimize the Chamfer distance between the input and output point clouds. Parameters were updated with the Adam optimizer; the learning rate started at 4 × 10^−2^ and was decayed to 1 × 10^−7^ over the course of training. Mini-batch size was 32. Training proceeded through 60 sequential runs: each run continued from the model state at the end of the previous run, with the learning rate held constant within a run and stepped down across consecutive runs to realize the overall decay schedule. In each run, objects were randomly split into 80% training and 20% validation partitions; the validation partition was used only to monitor training behavior and did not drive early stopping or other training decisions. A fixed random seed (1) was used throughout. The total training spanned 6,737 epochs across the 60 runs, and the model from the end of the 60th run was used for all subsequent analyses.

For the diffusion-based autoencoder, the model was trained end-to-end by minimizing the DDPM noise-prediction objective, computed as the mean squared error over all points and coordinate dimensions. Parameters were updated with the Adam optimizer using the default momentum parameters (*β*_1_ = 0.9, *β*_2_ = 0.999) and no weight decay; the learning rate was initialized at 1 × 10^−3^ and linearly decayed to 1 × 10^−4^ between iterations 150,000 and 300,000, and fixed thereafter. Gradients were clipped to a maximum norm of 10. Mini-batch size was 128, and a fixed random seed (2020) was used throughout. Objects were randomly split into 90% training and 10% validation partitions, with the validation partition used only to monitor training behavior. Training was run for 1,000,000 iterations without early stopping, and the final model was taken from the checkpoint at 1,000,000 iterations.

### Target features for decoding

To extract the target 3D features for brain decoding, we converted each mesh into a point cloud by randomly sampling 2,500 points on the surface, which was fed into the 3D encoder. The input tensor was passed through the encoder of each autoencoder, and the output latent features were used as decoding targets. For AtlasNet, we extracted the unit activations after the final fully connected layer and batch normalization, but before the ReLU activation function, yielding a 1,024-dimensional feature vector. For the diffusion-based autoencoder, whose encoder does not include a ReLU activation at the output, the 256-dimensional feature vector was used as is.

All point clouds in the brain-decoding analyses (target features for decoding, predicted point clouds, and the candidate point clouds used for evaluation) were kept at their original size in a viewer-centered coordinate frame; no centering, scaling, or recentering was applied to the point clouds at feature extraction, reconstruction, or evaluation, so that the pipeline reconstructs the absolute size and the viewer-relative depth (camera-to-object distance) as well as the shape of each object. Point-cloud normalization was used only during autoencoder pretraining (unit-sphere normalization in AtlasNet’s default training pipeline; standardization to zero mean and unit standard deviation for the diffusion-based autoencoder; see Model training); it was not applied to the test stimuli, decoded outputs, or the Chamfer distance computation. The latent-feature standardization that takes place inside the brain decoder (3D feature decoding from brain activity, below) is a separate, scalar-level operation and does not act on the point-cloud geometry.

### Representational similarity analysis (RSA)

To characterize the structure of the latent space relative to image-, semantic-, and shape-based references, we performed representational similarity analysis (RSA)^51^. For each modality, we computed a representational dissimilarity matrix (RDM) from pairwise distances among 2000 training objects. For latent features and 2D rendered images, object-to-object distances were defined using pattern correlation distance (*d* = 1 − *r*), where *r* denotes the Pearson correlation. For 3D point clouds, we used the Chamfer distance. Semantic categories were defined using ShapeNet synsets, and semantic distances were derived from WordNet-based Wu–Palmer similarity^52^ by converting similarity to distance *d* = 1 − sim_WP_. To stabilize the computation, synsets in a direct parent–child relation in WordNet were merged into the parent synset. In addition, because objects in the animal category were collected from outside ShapeNet, we assigned them the synset “animal.n.01”. To quantify similarity between RDMs, we vectorized the upper-triangular entries of each RDM (excluding the diagonal) and computed Spearman’s rank correlation between the resulting vectors. RSA results are shown in Figure S2 (AtlasNet) and Figure S3D (diffusion-based autoencoder).

### 3D feature decoding from brain activity

Following the brain decoding framework in a previous study^24^, we trained feature decoders using 6,000 samples from the training dataset for each brain region. The decoder predicted activation of each unit in the target 3D features from a multi-voxel fMRI pattern. The training process began with voxel selection based on the correlation coefficient with the activation of each target unit using the training stimuli. 5,000 voxels with the largest absolute correlations in each ROI were selected as input to the decoders. The responses of the selected voxels were normalized using the mean and standard deviation computed from the training samples. Likewise, target unit activation was normalized using the mean and standard deviation computed from the training samples. For decoding, we used an L2-regularized linear regression model to predict normalized unit activation from the normalized multi-voxel fMRI patterns. The regularization parameter was fixed at 5,000 for all models, regardless of ROI or subject. We independently trained a decoder for each unit in the target feature (1,024 units for AtlasNet; 256 units for the diffusion-based autoencoder).

In the testing phase, each fMRI sample from the test dataset was normalized using the mean and standard deviation obtained from the training dataset, and the trained decoders were applied to these normalized samples to predict 3D features. For the main analyses, the test fMRI samples were trial-averaged across the eight repetitions of each stimulus before decoding; for the supplementary single-trial analysis (Figure S7B), each trial was decoded independently without averaging. The decoded features were then denormalized using the mean and standard deviation for each feature derived from the training dataset.

### Cross-validated decoding for rescaling parameters

To support the post-hoc rescaling step described below, we additionally performed cross-validated feature decoding using only training samples to estimate the standard deviation of decoded features. We split 120 runs of the training sample into training (117 runs) and test (3 runs) sets in which the test set included unique 50 stimuli not overlapping with the training set. The decoders were trained using the training set and then predicted features using the test set in the same manner as the training and test using the full dataset. We repeated this training–test split to cover all training samples and averaged the three held-out repetitions of each stimulus, yielding 2,000 stimulus-level cross-validated decoded features (one per training stimulus).

#### Decoded feature rescaling

Regularized regression shrinks decoded feature values toward the training mean, reducing the variance of decoded features relative to that of the ground-truth features. Because the autoencoder’s generator was trained on features at their original scale, this shrinkage degrades the input distribution that the generator expects. We therefore rescaled the decoded features before passing them to the generator. The scaling parameters were estimated solely from training-set decoded features (no test data) as

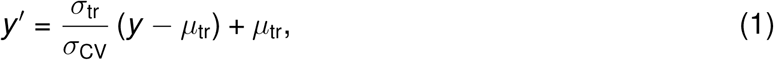

where *y* is a decoded unit activation, y ′ is the rescaled decoded unit activation, *µ*_tr_ and *σ*_tr_ are the mean and standard deviation of the ground-truth target-feature unit activations in the training samples (i.e., the target 3D features, not brain-decoded values), and *σ*_CV_ is the standard deviation of the corresponding cross-validated brain-decoded features. For AtlasNet, a ReLU operation was then applied to the rescaled features to match the activation function at the encoder output, bringing the decoded features—whose decoding target was the pre-ReLU activation—back into the same post-ReLU embedding space as the encoder output *z* that the generator takes as input. For the diffusion-based autoencoder, whose encoder does not include an output ReLU, no such operation was applied. The output of this step—a vector of rescaled (and, for AtlasNet, ReLU-clipped) decoded features—is the input to the 3D reconstruction stage described next.

#### Decoding evaluation

We evaluated decoding performance of 3D feature prediction with two complementary metrics: (1) profile correlation, defined as the Pearson’s correlation coefficient between the true and decoded features across test stimuli for each unit, and (2) pairwise identification accuracy, applied at the feature level. The pairwise identification procedure is defined in Evaluation of 3D reconstruction below; for feature-level decoding it is applied with Pearson’s correlation between the decoded and candidate latent features—a pattern correlation computed across feature units within each sample, as opposed to the profile correlation computed across stimuli for each unit—in place of the Chamfer distance between point clouds. The candidate pool consisted of the 80 test natural objects for natural-object stimuli and the 30 artificial objects for artificial-object stimuli. Profile correlation quantifies the per-unit linear agreement between decoded and true features, while pairwise identification accuracy quantifies whether the pattern of the decoded features for each sample is closer to the ground-truth feature of the true 3D structure than to those of false candidates.

### 3D reconstruction

Given the rescaled decoded features (3D feature decoding from brain activity, above), the autoencoder’s generator was used to reconstruct an explicit 3D representation of the stimulus—first as a point cloud and, for AtlasNet, additionally as a triangle mesh. The generator was used as released: we did not retrain or fine-tune the generator weights at the brain-decoding stage, so all geometric structure in the output comes from (i) the pretrained generator and (ii) the rescaled decoded features.

### Point-cloud prediction

For AtlasNet, the generator deforms a fixed sphere template of *M* = 2,562 points conditioned on the decoded latent feature z′ ∈ R^1024^. A multi-layer perceptron maps each template point’s 2D parametric coordinates to a 3D position, with z′ concatenated to the input at every layer, yielding a predicted point cloud *P*_rec_ ∈ R*^M^*^×3^. Because every point in *P*_rec_ corresponds to a known location on the sphere template, the topological structure of the template is preserved in the prediction, which is exploited at the surface-generation step below.

For the diffusion-based autoencoder, the generator predicts the denoising vector at each of *T* = 200 steps with a linear variance schedule, conditioned on the decoded latent feature *z*^′^ ∈ R^256^. Starting from independent Gaussian noise samples for 2,500 points, the denoising network (a six-layer multi-layer perceptron) iteratively refines the point positions across all *T* steps, yielding the predicted point cloud. Unlike AtlasNet, this generator does not maintain a fixed point-wise correspondence to a template, so the predicted points are an unordered set.

### Surface generation

For AtlasNet, a triangle mesh was obtained from the predicted point cloud using the closed-form mesh generation function provided in the original implementation, which inherits the known triangulation of the sphere template and applies it to the deformed point positions. This template-based scheme avoids the failure modes of post-hoc surface reconstruction (e.g., spurious topology from noisy point clouds) and produces a watertight, well-defined mesh whenever the template-deformation step succeeds. For the diffusion-based autoencoder, the predicted point set has no such correspondence; we therefore did not attempt to generate a mesh and used the point cloud directly for all quantitative analyses.

The template-based mesh used here captures geometry only; it does not capture material, texture, or view-dependent appearance (see Discussion for prospective extensions using richer scene representations).

### Evaluation of 3D reconstruction

This subsection defines the metrics applied to the explicit 3D reconstructions output by the pipeline. The same pairwise identification logic also underlies the feature-level decoding evaluation in 3D feature decoding from brain activity, above.

#### Chamfer distance

We used the Chamfer distance (CD) between the two sets of point clouds as a metric of 3D geometric similarity. Given point clouds *P* ⊂ R^3^ and *Q* ⊂ R^3^, the Chamfer distance is defined as:

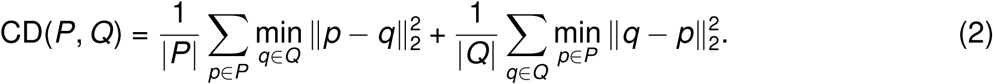

This metric measures the average squared distance from each point in one point cloud to its nearest neighbor in the other point cloud.

#### Identification accuracy

To evaluate reconstruction quality, we performed pairwise identification analysis based on Chamfer distance. The analysis quantified how well the true object can be identified among candidate objects given a reconstruction. For natural-object reconstructions, the 80 test natural objects served as candidates; for artificial-object reconstructions, the 30 artificial objects were used. For each reconstructed point cloud, we computed the Chamfer distance to the true point cloud of every candidate object (including that of the presented stimulus). Identification accuracy was defined as the proportion of false candidates for which the Chamfer distance to the true object was smaller than to the false candidate—that is, the reconstruction was geometrically more similar to the true object than to the alternative. For the same-category pairwise identification analysis (Figure S7A), false candidates were restricted to objects from the same category as the true object (chance = 50%) to test whether the pipeline distinguishes individual objects beyond category-level information.

#### Human evaluation

We performed behavioral experiments to assess the perceptual quality of the reconstructed 3D objects. This evaluation was conducted using reconstructions from the AtlasNet-based pipeline only. From each of the five subjects, we randomly selected 40 reconstructions of natural objects (20 from trained categories and 20 from novel categories) and all 60 reconstructions of artificial objects (30 from 2D rendered images and 30 from RDS reconstructions), resulting in 500 reconstructed objects in total, which were evaluated by seven evaluators. The evaluators were graduate students at Kyoto University who were not among the subjects of the fMRI experiments. We included reconstructions from the whole VC only.

Each reconstruction was evaluated in five pairwise identification trials. In each trial, the evaluators were presented with a reconstructed object along with two stimulus objects (Figure S7D): the true object and a false object. The reconstruction and both stimuli were shown rotating through 360^◦^ around the vertical axis over 4 s and repeated until response. Evaluators were required to select the stimulus (left or right) more similar to the reconstruction as a forced pairwise judgment (no “don’t know” option); response was disabled during the first 4-s rotation and unrestricted thereafter. Evaluators were instructed to judge the similarity by considering the overall 3D structures as well as the orientation and tilt.

For natural objects, the five trials for each reconstruction included one within-category false object and four between-category false objects; between-category false objects were randomly drawn from a different category. For artificial objects, the false objects were from the set of artificial objects in all trials. The left/right placement of the true and false stimuli was independently randomized on each trial, and the order of trials was randomized within each evaluator without separation by stimulus type. In total, the experiment comprised 2,500 trials, distributed across the seven evaluators: 200 trials were inherited from an initial pilot in which one evaluator assessed a subset of one subject’s reconstructions, and the remaining 2,300 trials were spread across the other six evaluators (each contributing 300 or 400 trials).

#### Principal axes in 3D bars

For the contour-matched RDS experiment, the orientation of each bar was summarized by the orientation of its longitudinal axis. We applied principal component analysis (PCA) to the 3D point cloud and defined the principal axis of the bar as the line of the first principal component. PCA was computed in 3D first; the resulting principal-axis direction was then projected onto the appropriate 2D plane (the top-view, *XY*, plane for horizontal bars; the side-view, *YZ*, plane for vertical bars) to yield a slant angle. Because the first principal component defines a line whose sign is arbitrary, we fixed the sign of the slant angle by the depth ordering of the bar ends: for horizontal bars, positive angles denote the right end nearer to and the left end farther from the observer (negative angles the reverse); for vertical bars, positive angles denote the lower end nearer and the upper end farther (negative angles the reverse). Principal axes were computed in this way for both the true and reconstructed slant bars. The angles of these axes, *θ*_stim_ (stimulus; the actual geometric slant recomputed for the presentation geometry, as described under Contour-matched RDSs) and *θ*_pred_ (reconstruction), serve as the variables of interest for the slant-prediction models in Statistical analyses below.

### Statistical analyses

#### Design and per-subject reporting

The primary unit of analysis is the individual subject. Following an intensive within-subject paradigm—widely advocated in cognitive neuroscience and central to our prior work—each subject contributes dense per-subject data sufficient for per-subject inference^13,14,24,42,43,45,46^. All experimental protocols and analysis pipelines were finalized through preliminary experiments before the main study began; subjects S1–S5 were then scanned under this fixed protocol, so all five datasets serve as confirmatory tests of the predefined design. Per-subject identification accuracy is computed by averaging across test stimuli within each condition and ROI (chance = 50%), shown as a single data point per subject, with its 95% confidence interval, in Figures 3F, 4F, 6D, and 6E. Within each subject, we tested whether identification accuracy exceeded chance (50%) with a one-sample *t* test applied to the per-stimulus accuracies.

#### Sample size

We studied five subjects (S1–S5); each is treated as an independent replication, so the primary sample sizes for inference are the numbers of test reconstructions and trials, not the number of subjects. The numbers of test stimuli were fixed before data collection at 40, 40, and 30 (trained-category natural, novel-category natural, and artificial objects, as 2D rendered images), exceeding the minimum required to detect a medium effect size (Cohen’s *d* = 0.5) with a one-sided *t* test at *α* = 0.05 (27 stimuli). The 30 artificial objects were additionally presented as RDSs, and the contour-matched RDS experiment used slant bars spanning eight nominal slant angles (Methods, Experimental design).

#### Group-level inference

To complement the per-subject statistics, cross-subject contrasts (condition or ROI comparisons) were assessed with linear mixed-effects models (LMMs) that include subject as a random effect. The main inference is frequentist (lmerTest in R; restricted maximum likelihood; Kenward–Roger small-sample correction; *α* = 0.05), with one-sided *p* values for directional hypotheses and two-sided otherwise. A parallel Bayesian LMM (Bambi/PyMC, weakly informative priors, NUTS sampler, four chains) serves as a robustness check, reporting the 95% highest-density interval (HDI) and the posterior probability of the hypothesised direction (Pr(dir.)). Confidence intervals and HDIs are 95% throughout. Key contrasts are reported inline in the Results, with the tables as the canonical record.

#### Whole-VC and sub-ROI models

Because the whole VC contains the four sub-ROIs (early VC, MT & neighbors, dorsal VC, ventral VC) together with DVT/ProS/POS1/POS2 (Table 1), we did not enter the whole VC and the sub-ROIs into one model. Region comparisons use two LMM forms, specified per analysis below: a *Whole VC model* (the whole VC treated as one region) and a *joint 4-ROI model* (the four sub-ROIs together, ROI as a fixed factor with early VC as baseline). Whole-VC estimates are reported in Tables S1 (frequentist) and S2 (Bayesian); sub-ROI estimates in Tables S3 (identification) and S4 (slant slope) (frequentist 95% CI and *p* value with Bayesian Pr(dir.)), with per-ROI slant slopes also shown as forest plots in Figure 7B.

#### Identification accuracy: RDS vs. 2D rendered images

The Whole VC and joint 4-ROI models are

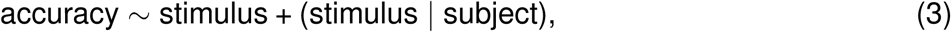

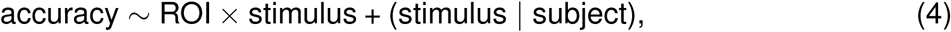

where stimulus is RDS vs. 2D rendered images (2D rendered as baseline). We use Wilkinson notation throughout: an intercept is implicit in the fixed- and random-effects parts, the operator × denotes full crossing (the crossed main effects together with their interaction), and a term such as (stimulus | subject) specifies a by-subject random intercept and random slopes for the listed terms. To avoid over-parameterization with only five subjects, the random effects were kept minimal: a by-subject intercept and a random slope for the within-subject predictor (stimulus, or *θ*_stim_ below) only, with no random slopes for ROI or its interactions. The inferential tests are the Whole-VC stimulus main effect (Tables S1, S2) and the ROI × stimulus interaction contrasts relative to early VC (Table S3). Per-ROI identification accuracies are shown in Figure 6E (AtlasNet) and Figure S4D (diffusion-based autoencoder); the within-ROI 2D-rendered-vs-RDS contrasts marked there are per-ROI simple effects from the joint 4-ROI LMM (estimated marginal means, Kenward–Roger), with the regional inference based on the ROI × stimulus interaction.

#### Slant prediction in contour-matched RDSs

The variables *θ*_stim_ and *θ*_pred_ are defined in Principal axes in 3D bars above. We tested whether reconstructed slant tracks stimulus slant across subjects, using the same two model forms as for identification accuracy—a Whole VC model and a joint 4-ROI model:

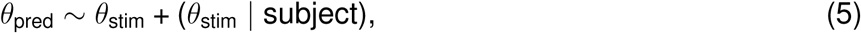

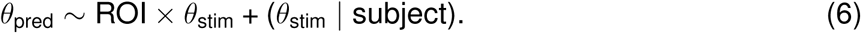

The Whole-VC slope is tested one-sided (slope *>* 0) and reported in Tables S1 (frequentist) and S2 (Bayesian). For the joint 4-ROI model, the regional inferential test is the ROI × *θ*_stim_ interaction relative to early VC (Table S4), exactly paralleling the identification ROI × stimulus interaction; the within-ROI *θ*_stim_ simple slopes (estimated marginal means, Kenward–Roger; each tested one-sided, slope *>* 0) are shown as forest plots in Figure 7B and tabulated in Table S4. These per-ROI simple slopes are themselves inferential within-ROI tests (slope *>* 0); the regional inference—whether a region’s slope differs from early VC—is carried by the interaction.

As a post-hoc, exploratory analysis, we also compared the slopes of the horizontal and vertical thin bars, using the matched runs of the second run type in which both orientations were presented together (a horizontal thin bar serving as a positive control). We added an orientation factor (horizontal vs. vertical) interacting with *θ*_stim_ in the Whole VC and joint 4-ROI models; the horizontal−vertical slope difference reported in Tables S1 and S2 is the *θ*_stim_ × orientation interaction (one-sided, horizontal *>* vertical).

## Supplementary Information

**Figure S1.**
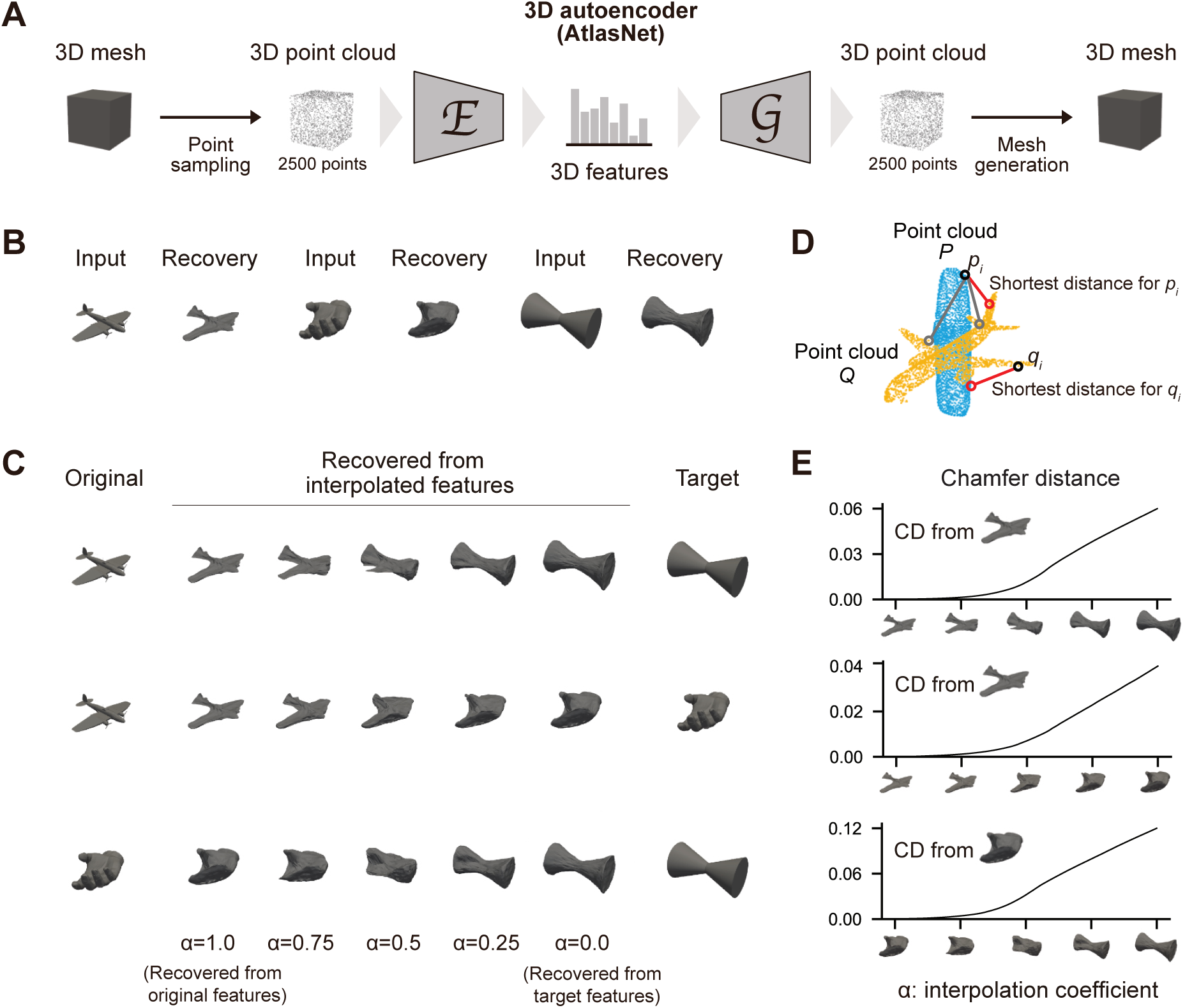
Latent-space recovery and local structure (AtlasNet). (A) Each 3D object is converted from a mesh into a point cloud (e.g., 2500 points) and passed through the 3D encoder *E* and generator *G*, yielding a recovered point cloud and mesh. (B) Recovery of an input object from its own latent feature, confirming shape preservation in latent space. (C) Linear interpolation between two objects’ latent features, *z*(*α*) = *αz*_1_ + (1 − *α*)*z*_2_, yields recovered point clouds that morph smoothly between endpoints; *α* is the interpolation weight for the left endpoint. (D) Chamfer distance (CD): mean bidirectional nearest-neighbor squared distance between two point clouds, capturing global geometric similarity. (E) Chamfer distance between the recovered point cloud at *α* = 1.0 and recovered point clouds from interpolated features increases monotonically as *α* is reduced from 1.0, indicating continuous shape changes with local latent-space movement.

**Figure S2.**
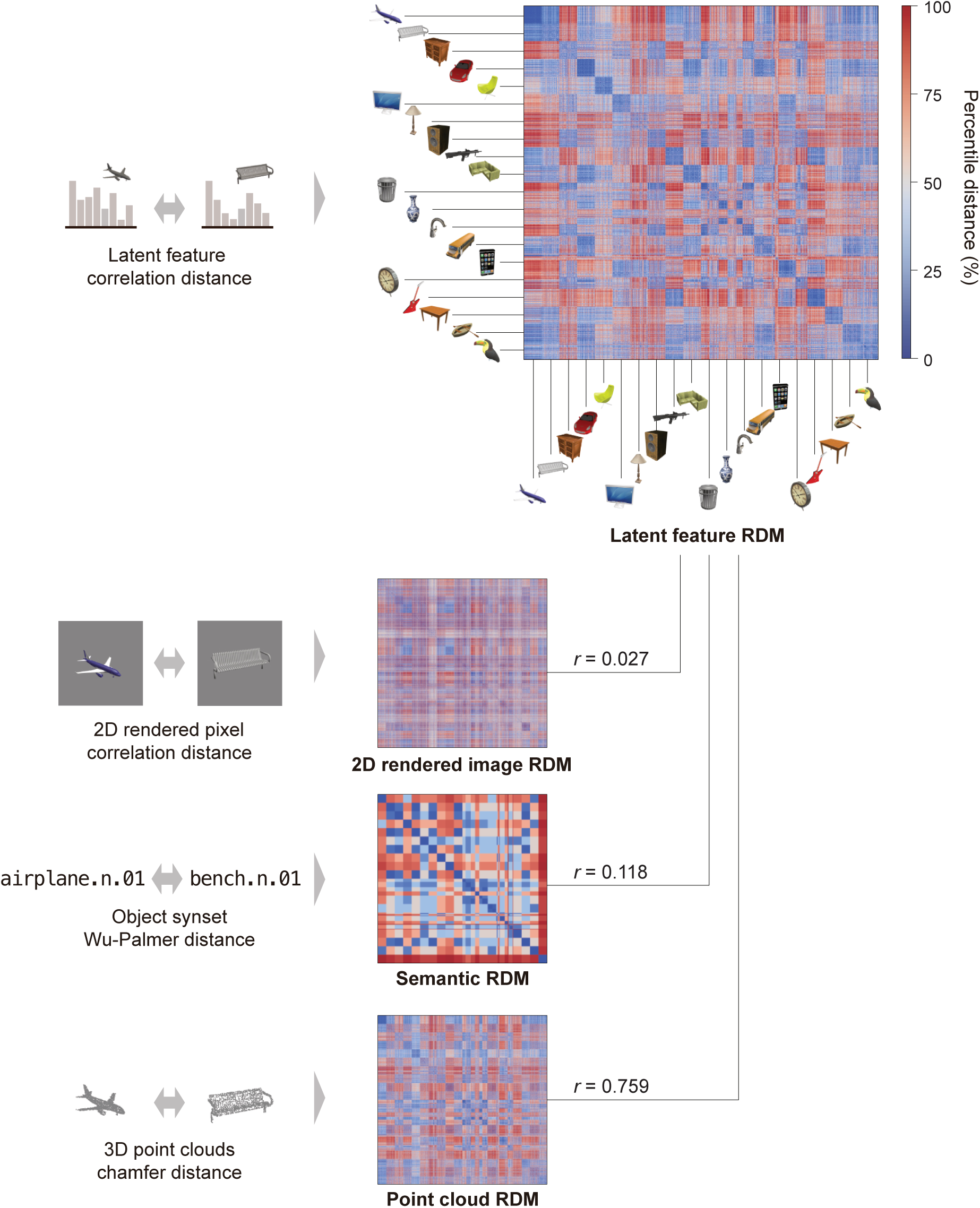
Representational similarity of AtlasNet latent features. Representational dissimilarity matrix (RDM) of AtlasNet latent features (top right) compared with RDMs from 2D rendered images, semantic categories (ShapeNet synsets with Wu–Palmer distance), and 3D point clouds (Chamfer distance). Spearman correlations between the latent-feature RDM and the three reference RDMs were 0.027, 0.118, and 0.759, respectively, indicating strongest alignment with point-cloud distances rather than pixel or semantic distances. Alternative semantic distances (path similarity and BERT-gloss embedding) gave comparable correlations (0.124 and 0.196). See Methods, Representational similarity analysis (RSA).

**Figure S3.**
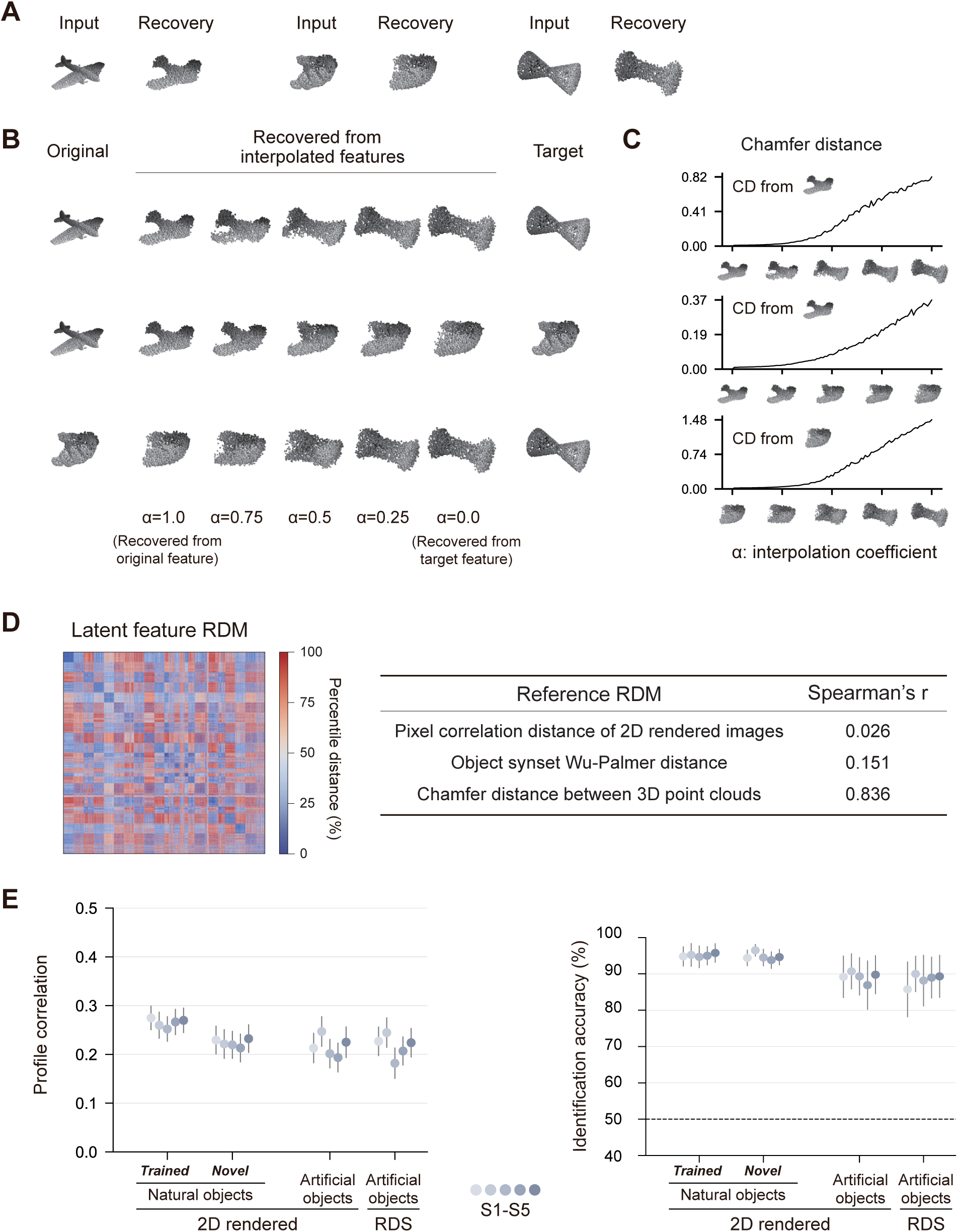
Diffusion-autoencoder latent features. (A–C) Recovery and latent-space interpolation, formatted as in Figure S1. (D) Representational similarity analysis. Spearman correlations between the latent-feature RDM (left) and reference RDMs (pixel correlation distance of 2D rendered images, Wu–Palmer synset distance, and Chamfer distance over 3D point clouds) were 0.026, 0.151, and 0.836, respectively. (E) 3D feature decoding using diffusion-based latent features: profile correlation between decoded and true features (left) and identification accuracy (right). Each dot is one subject (S1–S5); error bars are 95% CIs; dashed line indicates chance (50%).

**Figure S4.**
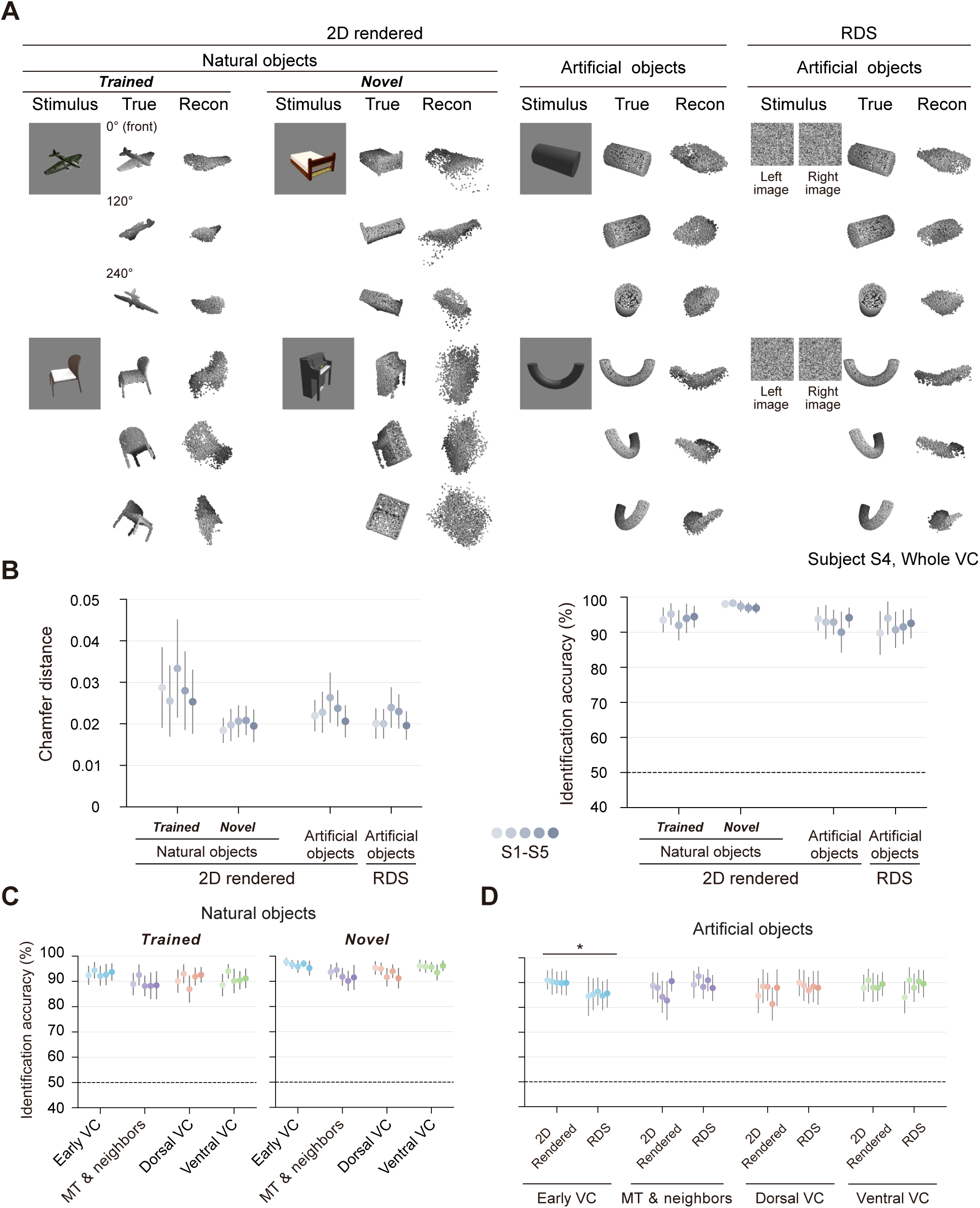
Diffusion-autoencoder reconstruction. (A) Example reconstructions. For each stimulus: visual stimulus, ground-truth 3D structure, and reconstructed 3D structure, each from three viewpoints. (B) Reconstruction performance: Chamfer distance between reconstructed and ground-truth point clouds (left) and identification accuracy based on Chamfer distance (right). (C) Identification accuracy across ROIs for natural objects (2D rendered images). (D) Identification accuracy across ROIs for artificial objects under 2D rendered vs. RDS conditions; asterisks mark within-ROI differences between the 2D rendered and RDS conditions (per-ROI simple effects from the joint 4-ROI LMM; ^∗^*p <* 0.05), reached here only in the early VC. In (B–D), each dot is one subject (S1–S5); error bars are 95% CIs; dashed line indicates chance (50%).

**Figure S5.**
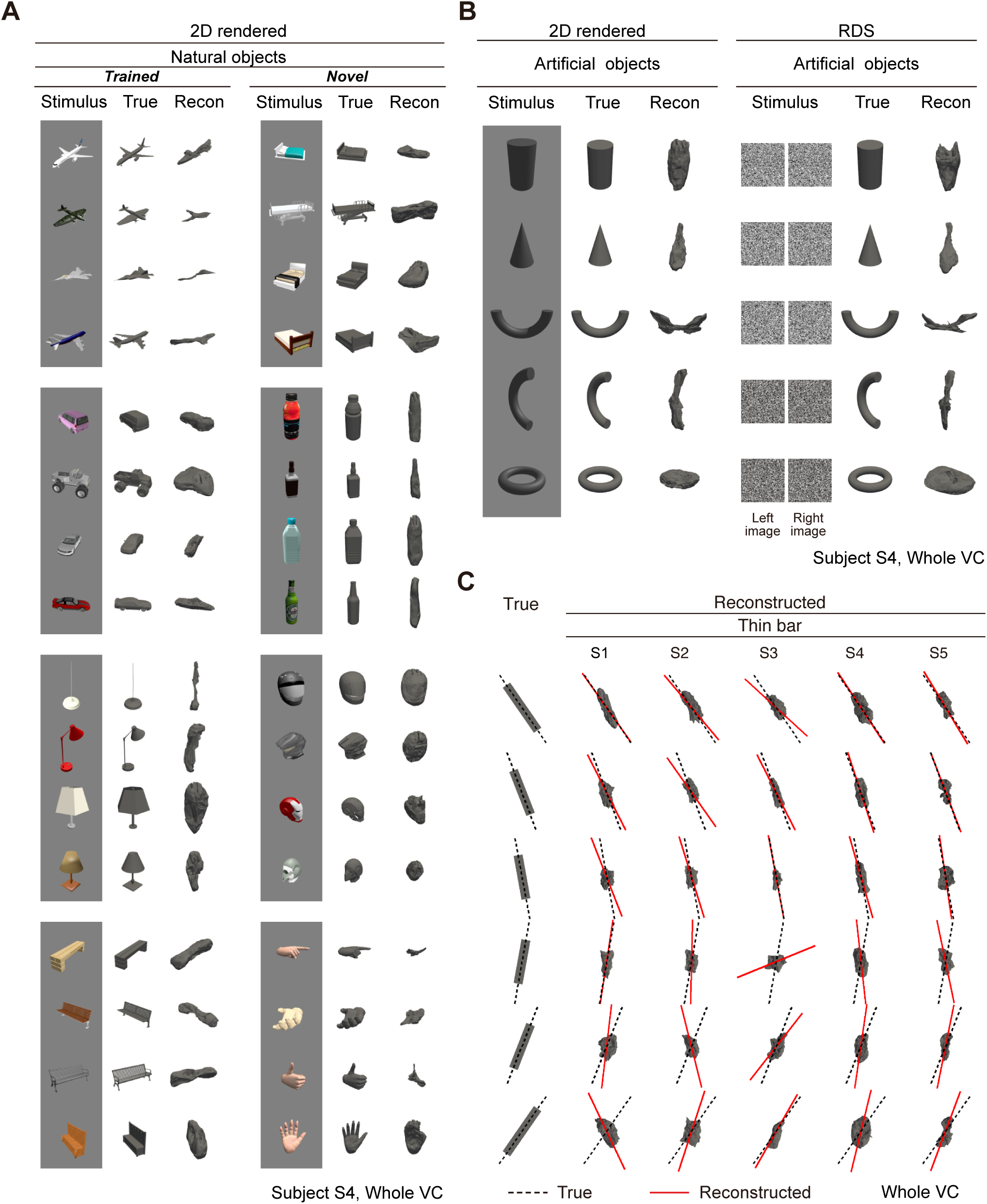
Additional reconstructions and cross-subject slant prediction. (A) Additional example reconstructions of natural objects (2D rendered images) for subject S4 (whole VC), grouped by trained and novel categories. (B) Additional example reconstructions of artificial objects (2D rendered images or RDSs) for subject S4 (whole VC). (C) Whole-VC reconstructions of contour-matched thin-bar RDSs across subjects (S1–S5); for each stimulus slant (rows), dashed black lines show the principal axis of the true point cloud and red lines that of the reconstruction.

**Figure S6.**
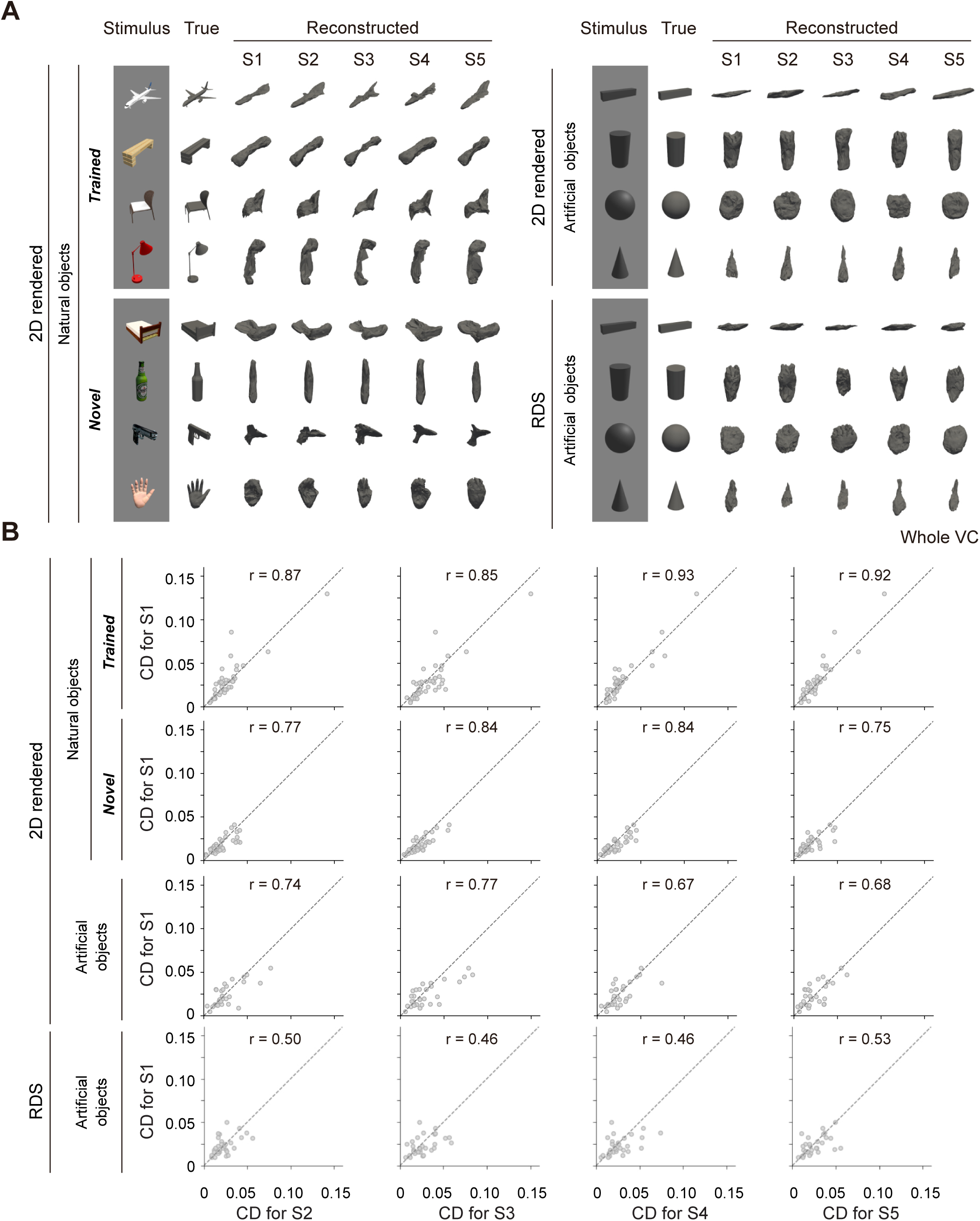
Similarity between subjects. (A) Qualitative cross-subject comparison. For each stimulus: visual stimulus, ground-truth 3D structure, and reconstructions from whole-VC activity for subjects S1–S5. (B) Quantitative cross-subject comparison. Scatter plots compare Chamfer distance (CD) for S1 with that of each other subject (S2–S5) on the same stimuli; rows are stimulus sets, columns are subjects. Each dot is one stimulus; Pearson’s correlation coefficient (*r*) is annotated in each panel; dashed diagonal indicates the identity line.

**Figure S7.**
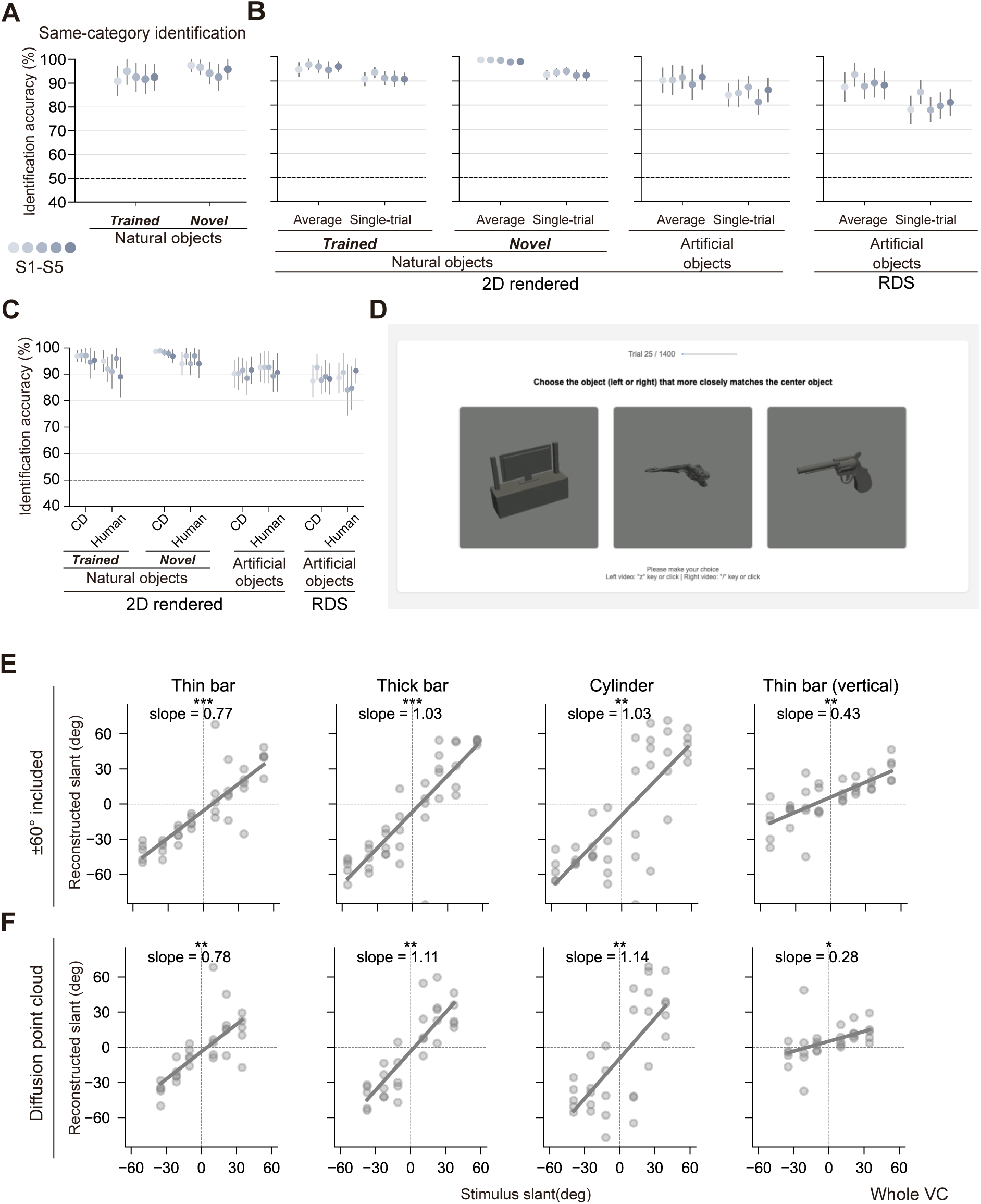
Reconstruction-evaluation controls and slant-prediction robustness. (A) Same-category identification accuracy for natural objects (false candidates restricted to the same category; chance = 50%). (B) Identification accuracy from single-trial vs. trial-averaged fMRI responses. (C) Identification accuracy based on Chamfer distance vs. human pairwise similarity judgments (AtlasNet, whole VC). (D) Human-evaluator pairwise similarity task interface. (E) Whole-VC slant regression including the largest-nominal-slant (±60^◦^ nominal) stimuli, for the thin bar, thick bar, cylinder, and vertical thin bar. (F) Corresponding whole-VC slant regression with the diffusion-based autoencoder. In (A–C), each dot is one subject (S1–S5); error bars are 95% CIs; dashed line indicates chance (50%). In (E) and (F), reconstructed slant (PCA-derived principal-axis angle) is plotted against stimulus slant; each dot is one subject at one stimulus slant angle; gray lines are fitted regression lines; asterisks indicate significant positive slopes (^∗^*p <* 0.05; ^∗∗^*p <* 0.01; ^∗∗∗^*p <* 0.001).

---

**Table S1.** Whole-VC frequentist statistics. Linear mixed-effects model estimates from the Whole VC models defined in Methods (Statistical analyses), fit to all five subjects (S1–S5). The identification accuracy contrast is two-sided; slant slopes are one-sided for the directional hypothesis (slope *>* 0). The horizontal–vertical slope difference is an exploratory, one-sided comparison (see Methods). Confidence intervals (CIs) are 95% throughout. The whole-VC slant slopes are plotted in Figure 5C, and the per-ROI counterparts in Figure 7B (Table S4); the Bayesian counterpart of this table is in Table S2. Slope differences (Δ*β*) are estimated within the Whole-VC orientation-interaction LMM and may not equal the arithmetic difference of the per-orientation slopes (see Methods).

| Contrast | $\beta$ | 95% CI | $p$ |
| --- | --- | --- | --- |
| Identification accuracy difference (RDS – 2D rendered) | –1.4% | [–6.2, +3.5] | 0.477 (two-sided) |
| Slant slope, thin bar (horizontal) | 0.772 | [0.275, 1.270] | 0.006 (one-sided) |
| Slant slope, thick bar | 1.113 | [0.644, 1.582] | 0.001 (one-sided) |
| Slant slope, cylinder | 1.154 | [0.392, 1.916] | 0.007 (one-sided) |
| Slant slope, thin bar (vertical) | 0.372 | [0.062, 0.683] | 0.015 (one-sided) |
| Slope difference, horizontal – vertical (exploratory) | 0.161 | [–0.077, 0.399] | 0.090 (one-sided) |

**Table S2.** Whole-VC Bayesian robustness. Bayesian linear mixed-effects model posterior summaries from the same Whole-VC models as in Table S1, fit with weakly informative priors (Bambi/PyMC, NUTS sampler, four chains). HDIs are 95% highest-density intervals. Pr(dir.) is the posterior probability that the effect lies in the hypothesised direction (positive for slant slopes and the slope difference; for the two-sided identification contrast, the posterior probability in the direction of the posterior median). Conclusions match the frequentist analysis (Table S1).

| Contrast | Post. median | 95% HDI | Pr(dir.) |
| --- | --- | --- | --- |
| Identification accuracy difference (RDS – 2D rendered) | –1.3% | [–5.6, +3.1] | 0.744 |
| Slant slope, thin bar (horizontal) | 0.764 | [0.275, 1.307] | 0.990 |
| Slant slope, thick bar | 1.106 | [0.590, 1.662] | 0.999 |
| Slant slope, cylinder | 1.151 | [0.284, 1.939] | 0.991 |
| Slant slope, thin bar (vertical) | 0.371 | [0.058, 0.716] | 0.981 |
| Slope difference, horizontal – vertical (exploratory) | 0.158 | [–0.075, 0.390] | 0.904 |

**Table S3.** Identification accuracy: per-ROI simple effects and ROI × stimulus interaction (joint 4-ROI LMM). All estimates come from the joint 4-ROI LMM defined in Methods (Statistical analyses). The left-hand columns report the per-ROI *simple effect* (the within-ROI RDS − 2D rendered difference at each region, estimated marginal means); the right-hand columns report the ROI × stimulus *interaction* relative to early VC (the regional difference from early VC, with early VC as reference). Both are inferential tests from the same model and both are two-sided; the simple effects are the values plotted in Figure 6E, whereas the regional inference is carried by the interaction. CIs are 95% (frequentist, Kenward–Roger); Pr(dir.) is the Bayesian posterior probability that the effect lies in the direction of the posterior median. Whole-VC values are in Tables S1 and S2. The slant-slope counterpart is Table S4.

| ROI | Per-ROI simple effect (within ROI) |  |  |  | Interaction vs. early VC |  |  |  |
| --- | --- | --- | --- | --- | --- | --- | --- | --- |
| | $\beta$ | 95% CI | $p$ | Pr(dir.) | $\Delta\beta$ | 95% CI | $p$ | Pr(dir.) |
| Early VC | –7.5% | [–12.3, –2.8] | 0.003 | 0.994 | (ref) | — | — | — |
| MT & neighbors | +3.1% | [–1.6, +7.9] | 0.185 | 0.894 | +10.7% | [+4.8, +16.5] | < 0.001 | 1.000 |
| Dorsal VC | +3.1% | [–1.7, +7.8] | 0.198 | 0.879 | +10.6% | [+4.7, +16.4] | < 0.001 | 1.000 |
| Ventral VC | –3.2% | [–7.9, +1.6] | 0.179 | 0.893 | +4.3% | [–1.5, +10.2] | 0.149 | 0.924 |

**Table S4.** Slant slope: per-ROI simple slopes and ROI × slant interaction (joint 4-ROI LMM). All estimates come from the joint 4-ROI LMM defined in Methods (Statistical analyses), separately for each bar type. The left-hand columns report the per-ROI *simple slope* (the within-ROI *θ*_stim_ slope at each region, estimated marginal means; one-sided test of slope *>* 0); the right-hand columns report the ROI × *θ*_stim_ *interaction* relative to early VC (the regional difference from early VC, two-sided, with early VC as reference). Both are inferential tests from the same model: the per-ROI simple slopes are the values plotted in Figure 7B, whereas the regional inference is carried by the interaction. CIs are 95% (frequentist, Kenward–Roger); Pr(dir.) is the Bayesian posterior probability that the effect lies in the hypothesised direction. The final block reports, as an exploratory analysis, the per-ROI horizontal−vertical thin-bar slope difference (the *θ*_stim_× orientation interaction within each ROI; one-sided, horizontal *>* vertical); its estimate, CI, *p*, and Pr(dir.) occupy the left-hand columns, and the interaction-versus-early-VC columns do not apply (“—”). Whole-VC values are in Tables S1 and S2. The identification counterpart is Table S3.

| ROI | Per-ROI simple slope (within ROI; slope $> 0$) | | | | Interaction vs. early VC | | | |
| --- | --- | --- | --- | --- | --- | --- | --- | --- |
| | $\beta$ | 95% CI | $p$ | Pr(dir.) | $\Delta\beta$ | 95% CI | $p$ | Pr(dir.) |
| *Thin bar (horizontal)* |  |  |  |  |  |  |  |  |
| Early VC | 0.291 | [−0.124, 0.707] | 0.074 | 0.889 | (ref) | — | — | — |
| MT & neighbors | 0.777 | [0.361, 1.192] | 0.001 | 0.993 | +0.485 | [0.127, 0.843] | 0.008 | 0.996 |
| Dorsal VC | 1.193 | [0.777, 1.608] | $< 0.001$ | 0.999 | +0.901 | [0.543, 1.259] | $< 0.001$ | 1.000 |
| Ventral VC | 0.612 | [0.196, 1.027] | 0.004 | 0.984 | +0.320 | [−0.038, 0.679] | 0.079 | 0.962 |
| *Thick bar* |  |  |  |  |  |  |  |  |
| Early VC | 0.640 | [0.109, 1.170] | 0.011 | 0.971 | (ref) | — | — | — |
| MT & neighbors | 0.526 | [−0.005, 1.057] | 0.026 | 0.950 | −0.114 | [−0.683, 0.456] | 0.693 | 0.660 |
| Dorsal VC | 1.150 | [0.619, 1.681] | $< 0.001$ | 0.996 | +0.510 | [−0.059, 1.080] | 0.078 | 0.962 |
| Ventral VC | 0.738 | [0.207, 1.269] | 0.005 | 0.980 | +0.099 | [−0.471, 0.668] | 0.732 | 0.633 |
| *Cylinder* |  |  |  |  |  |  |  |  |
| Early VC | 0.621 | [0.058, 1.183] | 0.016 | 0.963 | (ref) | — | — | — |
| MT & neighbors | 0.730 | [0.167, 1.292] | 0.007 | 0.979 | +0.109 | [−0.557, 0.775] | 0.747 | 0.630 |
| Dorsal VC | 1.380 | [0.818, 1.942] | $< 0.001$ | 0.997 | +0.759 | [0.093, 1.425] | 0.026 | 0.988 |
| Ventral VC | 0.507 | [−0.055, 1.070] | 0.037 | 0.937 | −0.113 | [−0.779, 0.553] | 0.736 | 0.631 |
| *Thin bar (vertical)* |  |  |  |  |  |  |  |  |
| Early VC | 0.062 | [−0.139, 0.263] | 0.260 | 0.706 | (ref) | — | — | — |
| MT & neighbors | 0.316 | [0.115, 0.517] | 0.002 | 0.987 | +0.254 | [0.046, 0.462] | 0.017 | 0.992 |
| Dorsal VC | 0.543 | [0.342, 0.743] | $< 0.001$ | 0.999 | +0.481 | [0.273, 0.689] | $< 0.001$ | 1.000 |
| Ventral VC | 0.139 | [−0.062, 0.340] | 0.080 | 0.884 | +0.077 | [−0.131, 0.285] | 0.465 | 0.764 |
| *Thin bar: horizontal – vertical slope difference (exploratory; one-sided, $H > V$)* |  |  |  |  |  |  |  |  |
| Early VC | 0.047 | [−0.180, 0.274] | 0.341 | 0.660 | — | — | — | — |
| MT & neighbors | 0.259 | [0.033, 0.486] | 0.013 | 0.986 | — | — | — | — |
| Dorsal VC | 0.142 | [−0.084, 0.369] | 0.108 | 0.892 | — | — | — | — |
| Ventral VC | 0.139 | [−0.087, 0.366] | 0.114 | 0.884 | — | — | — | — |

